# Tau-seed interactome analysis reveals distinct functional signatures in Alzheimer’s disease across model systems

**DOI:** 10.1101/2025.06.17.660179

**Authors:** Pablo Martinez, Henika Patel, Yanwen You, Daniella Lopes, Armando Amaro, Nur Jury-Garfe, Yuhao Min, Javier Redding-Ochoa, Sayan Dutta, Jean-Christophe Rochet, Nilüfer Ertekin-Taner, Juan C. Troncoso, Cristian A. Lasagna-Reeves

**Affiliations:** Department of Neurology, Baylor College of Medicine, Houston, TX, USA; Stark Neurosciences Research Institute, Indiana University School of Medicine, Indianapolis, IN, USA; Department of Neuroscience, Mayo Clinic, Jacksonville, FL, USA; Division of Neuropathology, Department of Pathology, Johns Hopkins University School of Medicine, Baltimore, MD, USA; Purdue Inst. for Integrative Neuroscience, Purdue University, Lafayette, IN, USA; Dep. of Medicinal Chemistry and Molecular Pharmacology, Purdue University, Lafayette, IN, USA; Department of Neurology, Mayo Clinic, Jacksonville, FL, USA

**Keywords:** Alzheimer’s disease, Tau, Tau-seed, Tauopathies, omics, interactome, screening, aggregation

## Abstract

Tau aggregates propagate through the brain in a prion-like manner in Alzheimer’s disease (AD) and other tauopathies, but the molecular identity and functional partners of the seeding-competent Tau species remain poorly defined. Here, we present an unbiased proteomic profiling of a high-molecular-weight (HMW) Tau-seed isolated from AD patient brains. We contrast this interactome with that of a biochemically similar, seeding-incompetent HMW-Tau species from age-matched healthy controls. Despite comprising less than 5% of total Tau in the brain, Tau-seed associates with a distinct set of proteins enriched in synaptic, mitochondrial, and vesicle-trafficking functions. Cross-species functional screening in *Drosophila* and mouse models identifies interactors that modulate Tau toxicity and seeding. Spatially resolved analysis of postmortem AD brains reveals heterogenous co-deposition of these proteins with Tau aggregates, suggesting functionally distinct Tau-seed complexes. Together, this dataset provides a framework for understanding selective Tau-seed toxicity and identifies candidate regulators of Tau propagation with therapeutic potential.

## INTRODUCTION

The pathological aggregation of the microtubule-associated protein Tau is a primary histopathological hallmark of Alzheimer’s disease (AD), progressive supranuclear palsy (PSP), and various neurodegenerative disorders collectively referred to as Tauopathies^1,2^. While the basic physiological role of Tau on microtubule dynamics is widely documented^3,4^, numerous interactome studies indicate that Tau directly interacts with proteins and complexes involved in diverse biological functions beyond those associated with microtubule stability^5–7^. Under pathological conditions, Tau undergoes misfolding and hyperphosphorylation, forming soluble high-molecular-weight (HMW) aberrant Tau species with seeding and propagating capabilities, termed “Tau-seeds”^8–11^. Recent evidence indicates that this HMW-Tau species is capable of suppressing complex-spike burst firing in hippocampal CA1 neurons in AS brains, linking them to the functional impairment of memory-related neuronal circuits independently of β-amyloid pathology^12^. Interestingly, similar HMW-Tau species have also been detected in brains of individuals with asymptomatic AD pathology (AsymAD); however, these forms lack seeding activity, suggesting that not all HMW-Tau species are pathogenic and may exhibit distinct structural or biochemical properties^13^. The formation of Tau-seeds may disrupt existing protein interactions or create new ones, resulting in detrimental changes within the brain network^14^. The study of the mechanisms behind Tau-seed formation, aggregation and propagation has gained significant research interest; however, the cellular processes and the specific modifiers of pathogenic Tau remain poorly understood. Many efforts have been made to understand these aberrant Tau interactions. For instance, different interaction studies indicate that direct Tau interaction with specific or larger complexes regulate Tau pathophysiology^15–17^. While recent studies have characterized total Tau interactors in *in vivo* tauopathy models^18–21^ and induced pluripotent stem cell-derived neurons^6^, no studies have determined the interactome of high-seeding-competent Tau species in AD and their role on aggregation and overall pathogenesis.

Previous work from our lab identified a form of soluble HMW-Tau with high seeding activity in the PS19 tauopathy mouse model. Isolated Tau-seed interactome analysis revealed that the presynaptic scaffolding protein Bassoon (BSN) is a critical interactor of the Tau-seed and exacerbates Tau-seeding and toxicity *in vitro* and *in vivo*. Moreover, BSN downregulation decreased Tau spreading and mitigated synaptic and behavioral deficits in this tauopathy model^11^. These findings highlight the critical role of Tau-seed interactors as potential pathogenic stabilizers, thereby enhancing aggregation and propagation.

Here, to overcome the limitations of prior interactome studies based on whole human brain lysates, which dilute disease-relevant signals by averaging across heterogeneous Tau species, we performed a proteomic analysis focused on a specific size-exclusion chromatography (SEC) fraction containing a defined HMW-Tau species, isolated using a total Tau antibody. We integrated quantitative proteomic analysis with functional screening and validation in *D. melanogaster* and a mouse model of tauopathy to characterize the human Tau-seed interactome isolated from AD and age-matched control brain cases. This integrated approach provides a systematic strategy for identifying and functionally validating Tau-seed interactors with a driving role in Tau pathology. Using unbiased quantitative mass spectrometry (MS), we identified proteins that aberrantly associate with AD-derived Tau-seeds and confirmed that only a small portion of total Tau forms a HMW complex with high seeding activity in AD patient brains. This seeding-competent HMW-Tau complex is enriched in synaptic, mitochondrial, and vesicle-trafficking proteins, suggesting that Tau aggregation and propagation may be intimately linked to key cellular pathways involved in neuronal communication and mitochondria dynamics. Furthermore, Weighted Gene Co-expression Network Analysis (WGCNA) of AD Tau-seed interactome revealed disease-specific reorganization of protein-protein interactions. To functionally assess the role of Tau-seed interactors in disease progression, we performed a large-scale genetic screen in a *Drosophila* tauopathy model, targeting 115 AD Tau-seed interactors. Our findings demonstrated that genetic downregulation of these interactors modulates Tau toxicity and aggregation, suggesting their direct involvement in Tau pathology. Integrative analysis with human post-mortem multi-omics datasets highlighted perturbations of these Tau-seed interactors. Moreover, follow-up validation studies in a tauopathy mouse model confirmed that downregulation of selected key Tau interactors modulates Tau-seeding activity, either attenuating or exacerbating it. Taken together, our findings suggest that the Tau-seed interactome plays an active and heterogenous role in forming pathogenic Tau species with high seeding activity in AD pathology. By identifying key molecular components involved in this process, our study provides critical insights into the mechanisms underlying Tau pathology and a resources of novel candidate targets for future therapies.

## RESULTS

### An HMW-Tau species is responsible for high seeding activity in AD brains

We previously developed a streamlined protocol to isolate and characterize Tau-seeds from a range of Tau models, including cell, mouse, *Drosophila*, and human brain samples^11,22^. To isolate and characterize Tau-seed species from AD and age-matched control brain samples, we performed SEC on Tris-buffered saline (TBS)-soluble extracts from the middle frontal gyrus (MFG) (Figure 1A, Figure S1A). Seeding activity was then assessed in each SEC fraction by transfecting them into Tau biosensor cells expressing the aggregation-prone repeat domain (RD) of Tau containing the P301S mutation and quantifying integrated FRET density by flow cytometry (Figure 1A and S1B). As we previously reported in a tauopathy mouse model^11^, the strongest seeding activity in AD brains was detected in fraction number 9 (SEC-F9) (Figure 1B, top panel), the void volume fraction containing high-molecular-weight (HMW) proteins larger than 2,000 kDa. This fraction contained less than 5% of total Tau, as quantified by enzyme-linked immunosorbent assay (ELISA) for total human Tau (Figure 1B, bottom panel). Interestingly, the age-matched controls had similar levels of Tau in SEC-F9 but lacked seeding activity, indicating the presence of an HMW-Tau species in control brains that differs from the AD Tau-seed in their pathogenic seeding properties (Figure 1B, bottom panel). To confirm that the Tau species in SEC-F9 were responsible for the observed seeding activity, Tau was depleted by immunoprecipitation (IP) using the HT7 antibody, resulting in a significant reduction of seeding activity in the flow-through (Figure 1C). Electron microscopy (EM) of Tau-IP material from SEC-F9 showed that the seed primarily consisted of twisted filament structures (Figure 1D and S1C), averaging 7 ± 0.13 nm in width (Figure 1E). No filamentous structures were detected in the immunoprecipitated Tau from control SEC-F9 (Figure S1C).

**Figure 1.**
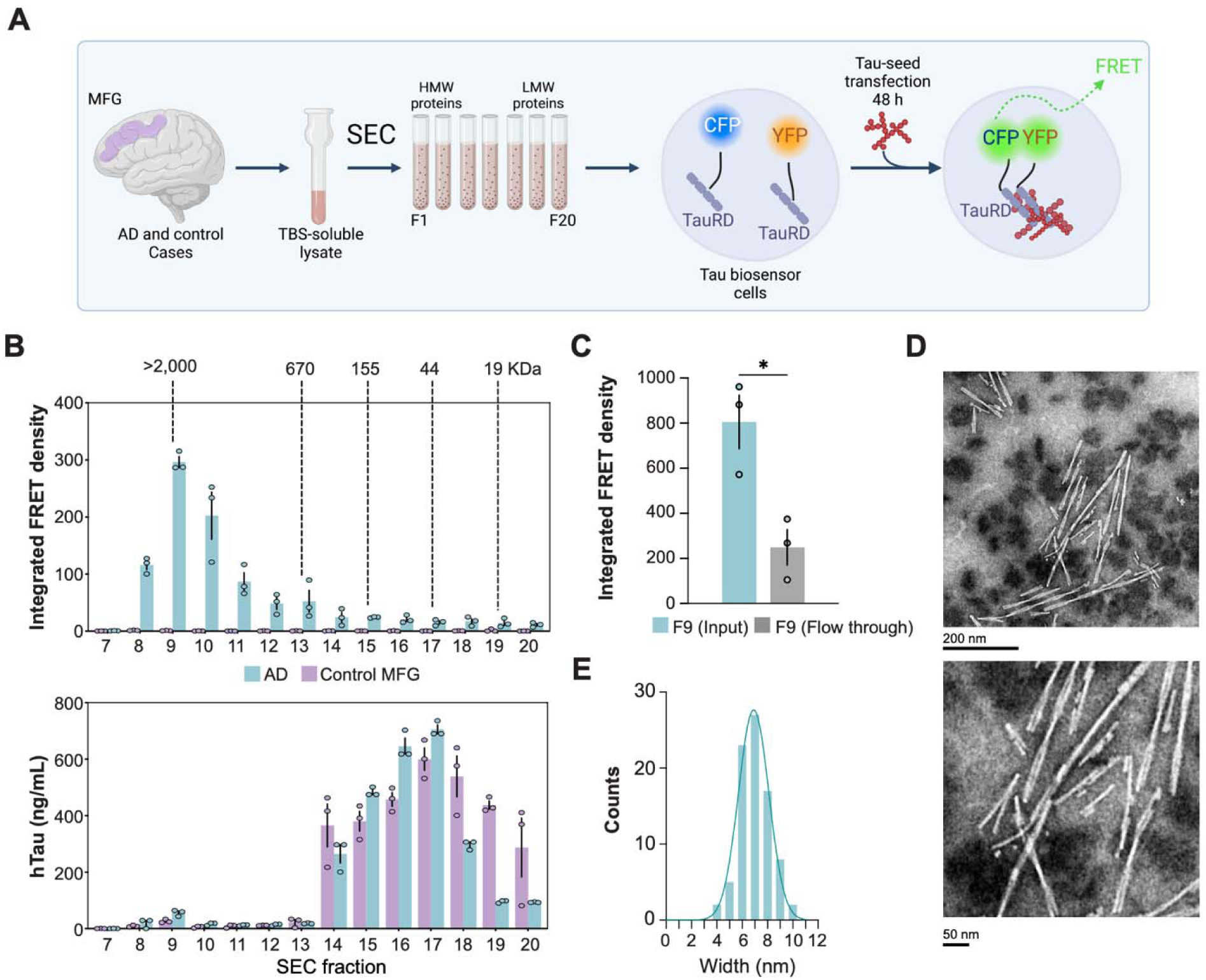
An HMW-Tau species drives high seeding activity in AD brains. (A) Schematic overview of the protocol used to assess seeding activity from SEC fractions of TBS-soluble brain lysates from AD and control cases. Created in BioRender. (B) Tau seeding activity (upper panel) indicating Tau seeding activity across SEC fractions, and corresponding quantification of total human Tau by ELISA (lower panel). Data are shown as the mean ± s.e.m (*n*=3). (C) Tau seeding activity of SEC fraction 9 (F9) from AD brain lysates before and after Tau immunoprecipitation (Tau-IP). Data are shown as the mean ± s.e.m., and significance was determined by unpaired two-tailed Student’s t-test (*p<0.05; *n*=3). (D) Electron micrographs showing twisted Tau filaments isolated from Tau-IP of AD F9 (scale bar, 200 nm; inset, 50 nm). (E) Width distribution of twisted Tau filaments shown in (D) (*n*=3).

### Proteomic analysis of AD Tau-seed and control HMW-Tau interactomes

To characterize the AD Tau-seed and control HMW-Tau interactomes, we immunoprecipitated Tau from SEC-F9 from AD and control brains and performed quantitative MS using TMT labeling. MS analysis identified 1,201 significant interactors in the AD Tau-seed F9 and 506 significant interactors in the control HMW-Tau fraction (Figure 2A-2C; Table S1). Gene Ontology (GO) enrichment analysis of cellular component (GO-CC) terms revealed that both interactomes localized to overlapping compartments, including the cytosol, cytoplasm, membrane, extracellular exosome, mitochondrion, focal adhesion, cytosolic ribosome, postsynaptic density, and glutamatergic synapse (Figure 2D and 2E). However, the AD Tau-seed interactome exhibited selective enrichment in axon-related compartments, suggesting that Tau-seed-interacting proteins are enriched in axonal regions (Figure 2D). Biological process (GO-BP) enrichment analysis identified shared enrichment in pathways related to translation and intracellular protein transport, indicating conserved functional processes between AD Tau-seed and control HMW-Tau interactomes (Figure 2F and 2G). Molecular function (GO-MF) enrichment analysis revealed distinct functional differences between the two interactomes. The AD Tau-seed interactome was enriched in structural constituents of the cytoskeleton and unfolded protein binding. Notably, structural constituents of the ribosome were also significantly enriched, suggesting that Tau-seed interactions with ribosomal proteins may contribute to proteostasis impairment, potentially through effects on ribosomal function^23^ (Figure 2H and 2I).

**Figure 2.**
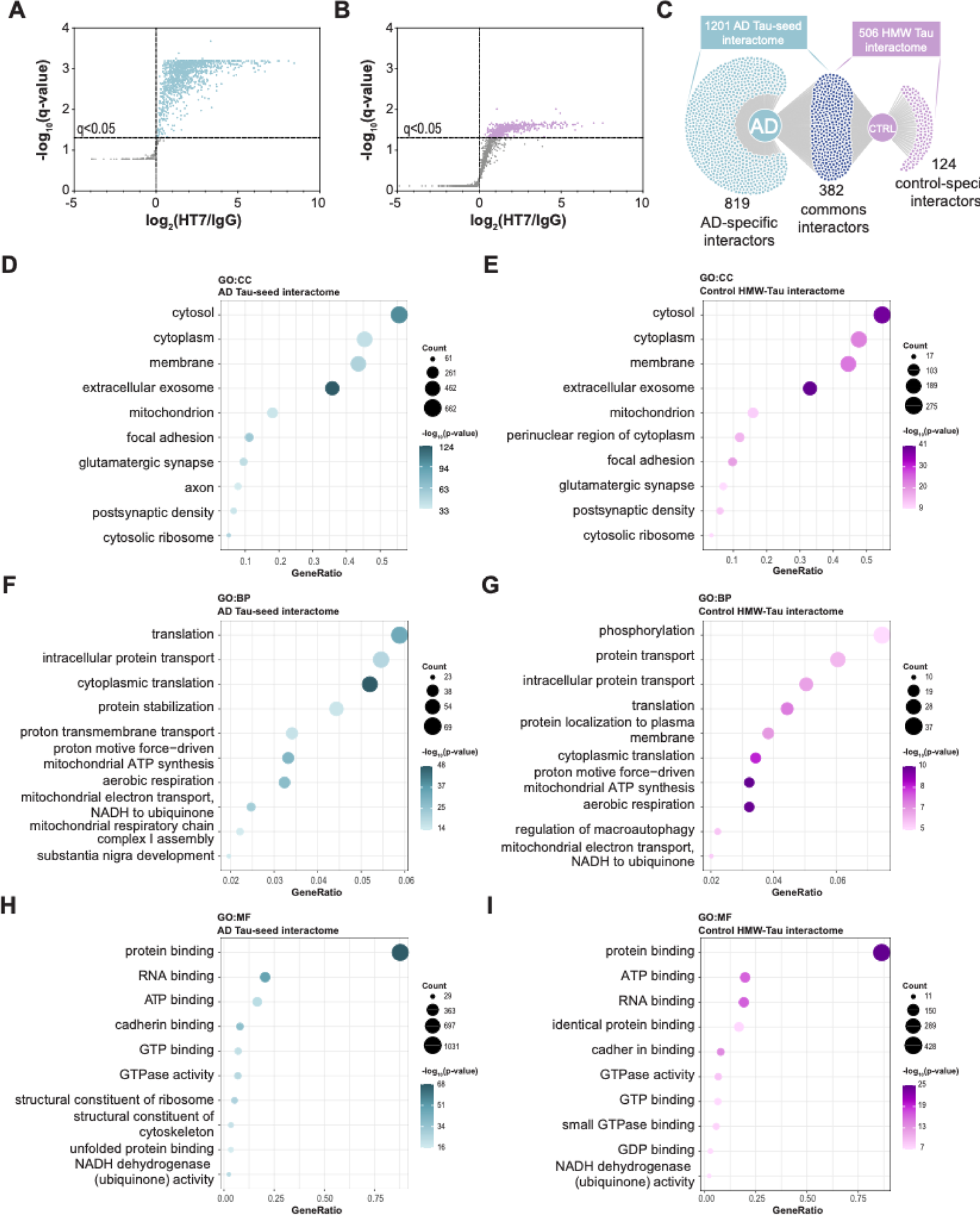
The AD Tau-seed interactome differs from control HMW-Tau (A and B) Volcano plots showing proteins enriched in immunoprecipitated Tau from AD Tau-seed showed in light green dost (A) and control HMW-Tau showed in light purple dots (B) identified by LC–MS/MS. See also Table S1. (C) DiVenn^74^ analysis displaying overlap between the AD Tau-seed, HMW-Tau, and shared interactomes. See also Table S1. (D–I) GO enrichment analysis of the AD Tau-seed and control HMW-Tau interactomes, showing terms enriched in the Cellular Component (D, E), Biological Process (F, G), and Molecular Function (H, I) categories. See also Table S1.

Common Tau interactors in AD and control brains may influence the formation of pathological Tau species through altered abundance levels. We identified 382 proteins shared between the AD Tau-seed and control HMW-Tau interactomes (Figures 2C and S2A). To explore the potential role of these common interactors in Tau pathology, we performed GO enrichment analysis on the shared protein set. GO-CC analysis highlighted broad localization across the cytosol, cytoplasm, membrane, and exosomes (Figure S2B), while GO-BP showed terms related to phosphorylation, protein transport, translation, membrane localization, and aerobic respiration (Figure S2C). GO-MF analysis revealed enrichment in ATP binding, protein binding, and RNA binding, highlighting their roles in molecular interactions and energy-related functions (Figure S2D). To assess disease-specific changes, we quantified the relative abundance of common interactors in AD vs. control brains, identifying 48 differentially abundant proteins (11 upregulated, 37 downregulated) (Figure S2E). Upregulated interactors were enriched in endocytic recycling pathways, suggesting increased endocytosis in AD, whereas downregulated interactors were linked to cellular biosynthesis, particularly translation, indicating impaired protein synthesis in AD (Figure S2F). These findings suggest that altered levels of shared Tau interactors may influence the formation and stability of pathogenic Tau species in AD.

To assess the phosphorylation profile of isolated AD Tau-seed and control HMW-Tau, we analyzed phosphorylation site occupancy using the MS described before. Although our MS approach was not specifically optimized for post-translational modification (PTM) quantification, it successfully detected several phosphorylated Tau residues. Specifically, phosphorylations at T181, T231, T235, and S262 were detected in the AD Tau-seed interactome, while pS404 was detected in both AD and control HMW-Tau interactomes (Figure S3A). No Tau phosphorylation sites were exclusively detected in the control HMW-Tau interactome (Figure S3A). To further validate the phosphorylation differences observed in AD Tau-seed, we performed a Meso Scale Discovery (MSD) biomarker assay to profile Tau phosphorylation in AD and control SEC fractions. Total Tau levels were evenly distributed across SEC fractions in both AD and control samples, consistent with our previous findings using ELISA assay (Figure S3B). Similarly, pT181 showed no significant differences in distribution between AD and control SEC fractions (Figure S3C). Notably, pT231 exhibited a distribution pattern that strongly resembled the distribution of seeding activity in AD, as highlighted by the gray overlay in the plot, suggesting a potential association between pT231 phosphorylation and Tau-seed pathogenicity (Figure S3D). Lastly, we also evaluated the distribution of pT217 due to its relevance as a biomarker in clinical studies^24–26^. Surprisingly, we did not find significant differences in pT217 levels between AD and control SEC fractions, including those HMW AD fractions with seeding activity (Figure S3E). Together, these findings reveal distinct phosphorylation signatures in AD Tau-seed compared to control HMW-Tau, highlighting a potential role of PTMs in modulating Tau-seeding activity and pathogenic propagation.

### Tau-seed and control HMW-Tau interactomes reveal distinct co-expression networks and functional pathways

Given the critical role of gene and protein coordination in biological function and the potential impact of network disruptions in AD pathology, we conducted a Weighted Gene Co-expression Network Analysis (WGCNA) as previously described^27^ to identify functional protein networks that could modulate Tau pathology. WGCNA categorized the AD Tau-seed interactome into 12 modules of highly co-abundant proteins (Figure 3A). The five largest modules were enriched in mitochondrial function (M1, 315 proteins), synaptic vesicle dynamics (M2, 253 proteins), intracellular protein trafficking (M3, 155 proteins), macromolecular disassembly regulation (M4, 154 proteins), and ribosome dynamics (M5, 149 proteins), representing the 85.4% of the total Tau-seed interactome (Figure 3A, Table S2). The HMW-Tau interactome from control brains produced 14 WGCNA modules, however, we did not find significantly enriched modules as found in the AD Tau-seed interactome (Figure 3B). Furthermore, module eigengene adjacency heatmap analysis revealed distinct module association patterns between AD and control groups (Figure 3C and 3D). Visualization of the protein correlation network revealed differences in the clustering and distribution of interactors within each Tau-seed module, compared to the HMW-Tau protein network (Figure 3E and 3F). This redistribution in the connectome is represented by changes in the module “neighborhood” and the distribution of the five largest modules in the Tau-seed interactome. Furthermore, the Tau-seed connectome contained groups of dense module clusters, suggesting a stronger correlation across several modules. Control HMW-Tau showed a homogeneous and less correlated network (Figure 3F), suggesting that the Tau-seed interaction network is reorganized in AD, potentially reflecting disease-specific alterations in key cellular processes and pathways, including mitochondrial function (module 1), and vesicle and synaptic dynamics (module 2). GO-BP enrichment analysis of the five largest AD Tau-seed modules revealed involvement in key biological processes, including mitochondrial respiration (M1), synaptic vesicle transport (M2), translation and post-synapse organization (M3), protein and RNA regulation (M4), and ribosome biogenesis (M5) (Figure 3G). Control HMW-Tau modules showed both similar and distinct annotations. Mitochondrial processes (M1) and synaptic vesicle transport (M2) overlapped with AD, while control-specific modules included nuclear protein localization (M3), calcium signaling (M4), and ER network organization (M5) (Figure 3H). Full GO enrichment analysis is available in Table S2.

**Figure 3.**
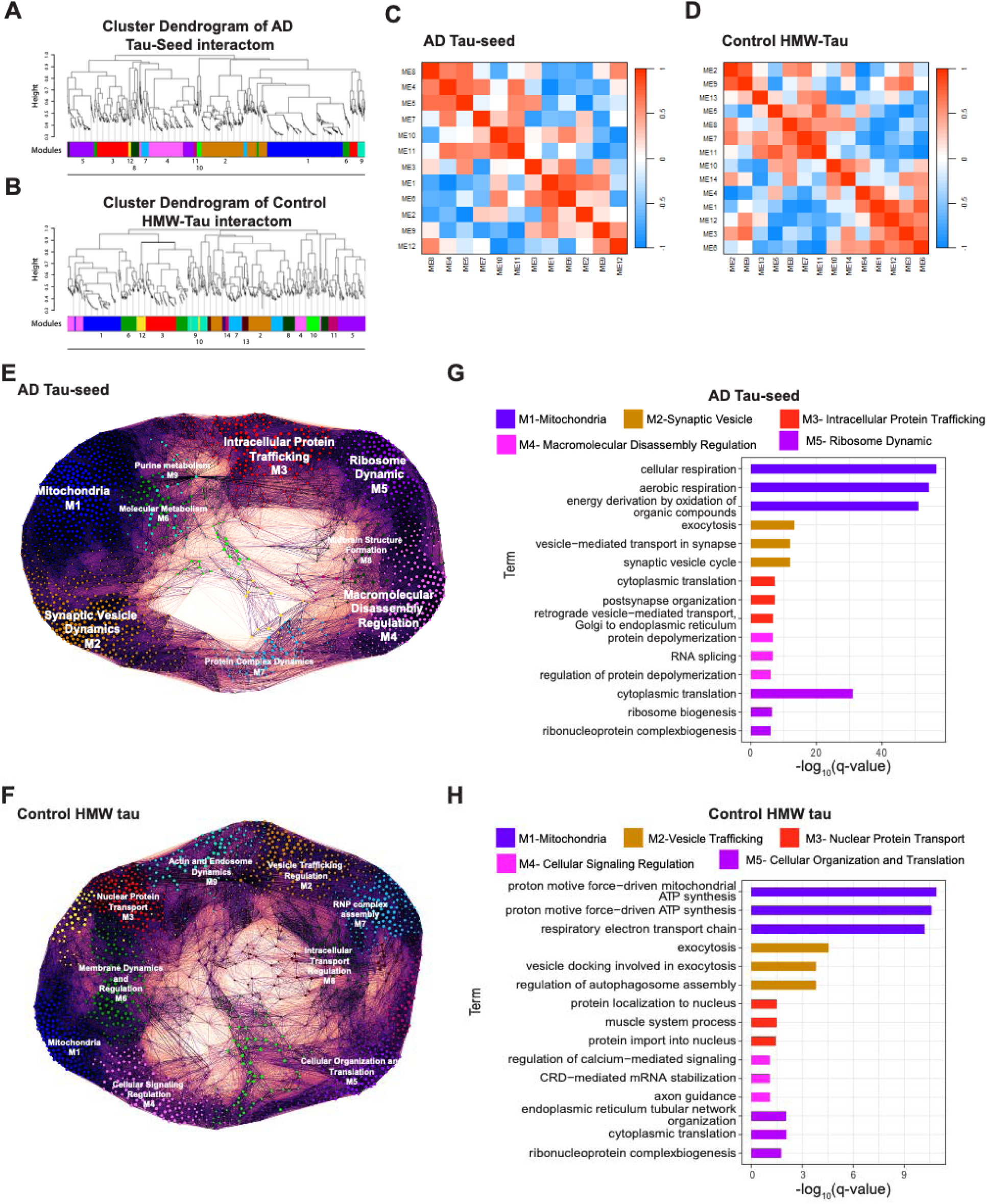
WGCNA analysis reveals distinct co-expression modules and functional profiles in AD Tau-seed (A and B) WGCNA clustering dendrograms for AD Tau-seed (A) and control HMW-Tau (B) interactomes derived from SEC fraction 9 Tau-IP. See also Table S2. (C and D) Module eigengene adjacency heatmaps showing inter-module correlation in AD Tau-seed (C) and control HMW-Tau (D). (E and F) Visualization of the protein correlation network of co-expression modules in AD Tau-seed (E) and control HMW-Tau (F). (G and H) GO enrichment of top five WGCNA modules from AD Tau-seed (G) and control HMW-Tau (H) interactomes (Biological Process category). See also Table S2.

### *Drosophila* Tau model-based screening identifies key Tau-seed modifiers

To evaluate the pathological relevance of Tau-seed interactors in AD, we compared the 1,028 proteins from the top five WGCNA modules derived from AD proteomic data with the Accelerating Medicines Partnership–Alzheimer’s Disease (AMP-AD) Agora list^28–30^. The Agora list is a curated resource of AD-associated genes and proteins identified through integrative multi-omic and systems biology approaches and is widely used to prioritize therapeutic targets in AD research^31^. This comparison revealed 180 overlapping proteins between the WGCNA-derived modules and the Agora list (Figure 4A). To functionally evaluate these candidates, we used a *Drosophila* tauopathy model that overexpresses human Tau^P301L^ under the GMR promoter, resulting in severe eye degeneration and recapitulating the Tau-seeding activity observed in PS19 mice and AD cases^11^. Among the 180 overlapping proteins, 142 (78%) had conserved *Drosophila* orthologs distributed across the top five WGCNA modules (M1: 54; M2: 34; M3: 16; M4: 22; M5: 16) (Figures 4B and 4C). Of these, 115 were available from the Bloomington *Drosophila* Stock Center (BDSC) for genetic screening (Figure 4A). Using RNA interference (RNAi) under the UAS/GAL4 system, we screened these interactors by evaluating Tau-seeding activity and eye degeneration phenotypes (Figure 4D). This analysis identified 29 Tau-seed interactors that significantly reduced eye degeneration, 2 that exacerbated it, 87 that decreased seeding activity, and 3 that enhanced it (Figure 4E). Notably, 25 interactors significantly reduced both eye degeneration and seeding activity (Figure 4E, Figure 4F and 4G; Figure S4A), all mapped across the top five WGCNA modules. STRING analysis of these interactors revealed synaptic-, mitochondrial-, and vesicle-related processes clusters with a known direct association to Tau (Figure 4H). Of these, 20 were uniquely enriched in the AD Tau-seed interactome, while 5 were shared with the control HMW-Tau interactome (Figure 4H).

**Figure 4.**
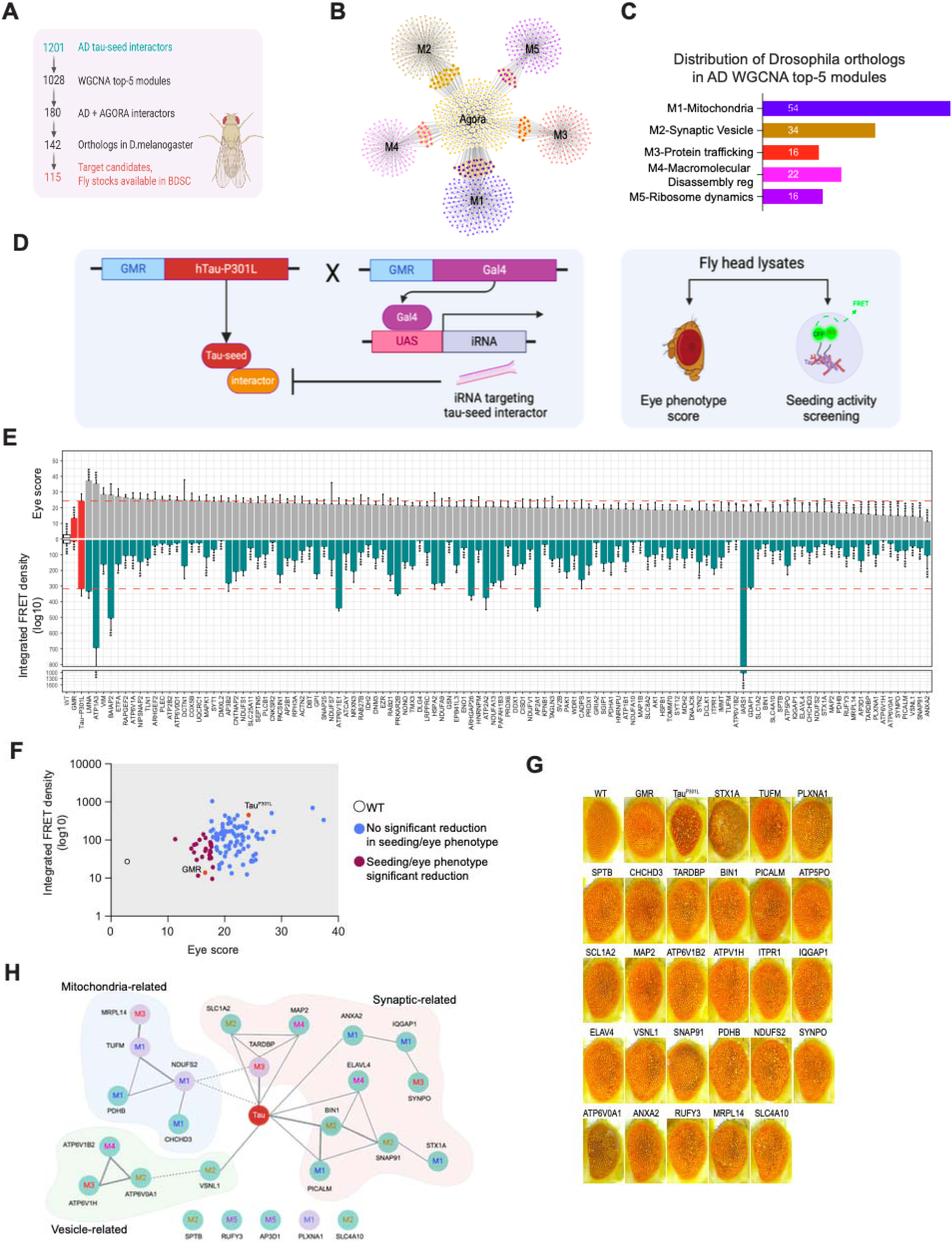
Identification of key Tau-seed interactors using a *Drosophila* tauopathy model (A) Workflow of candidate selection and *in vivo* screening in a *Drosophila* tauopathy model. Created in BioRender. (B) DiVenn analysis showing overlap between top WGCNA AD Tau-seed modules and the AMP-AD AGORA gene list. (C) Distribution of 115 *Drosophila* orthologs among AD Tau-seed WGCNA top modules. (D) Genetic strategy for RNAi-mediated knockdown in Tau^P301L^ flies^75^, used for eye phenotype and seeding assays. Created in BioRender. (E) Eye degeneration scores and seeding activity for 115 Tau-seed interactors. Red dotted line indicates the Tau^P301L^ tauopathy control. Data are presented as mean ± s.e.m. Statistical significance was determined by one-way ANOVA (*p<0.05, **p<0.005, *p<0.0005). *n* = 10 individual flies for eye score; for seeding activity, each point represents a pool of 20 fly heads collected from three independent eclosion events. See also Table S3. (F) Graphical representation of eye degeneration scores (X-axis) and seeding activity (Y-axis) for 115 Tau-seed interactors. Purple dots indicate the 25 downregulated proteins that significantly reduced both parameters; blue dots represent those with no significant effect for both measurements. WT is shown in white dot. GMR-negative and Tau^P301L^-positive controls are shown in red dots. (G) Representative eye phenotype images of 25 validated hits that reduced both eye degeneration and seeding activity, along with wild-type and control genotypes. See also Figure S4. (H) STRING network analysis of validated hits, annotated by WGCNA module; green ballons = AD Tau-seed-specific, purple ballons = shared with control.

### Multi-omic profiling links Tau-seed interactors to transcriptomic and proteomic signatures in AD

To understand the association of the 25 *Drosophila* hits in AD pathology, we performed GO enrichment analysis. We observed that these candidates are predominantly involved in synaptic vesicle dynamics, endo-lysosomal function, and Tau binding, validating the specificity of our approach. The enrichment in vesicle transport and lysosomal pathways suggest mechanisms by which Tau-seed interactors contribute to disease progression, potentially through impaired proteostasis and disrupted synaptic integrity (Figure 5A). To characterize the multi-omics changes of the 25 *Drosophila* hit genes in human brain, we retrieved mulit-omics datasets (transcriptomics and proteomics) across different brain regions. Specifically, we leveraged proteomic datasets from the Baltimore Longitudinal Study of Aging (BLSA)^32^ and Banner Sun Health Research Institute (Banner)^33^, totaling 245 samples (Table S3). We also used the RNAseq datasets generated by teams from Mayo Clinic (Mayo), Mount Sinai School of Medicine (MSSM), and Religious Orders Study/Memory and Aging Project (ROSMAP)^28–30,34^ (Table S3). We analyzed their association with AD diagnosis or Braak stage from seven transcriptomic datasets (Figure 5B), and proteomic data from the Banner and BLSA datasets (Figure 5C). Many of these hits are perturbed in AD brains and are significantly associated with AD diagnosis or Braak stage. Only two genes (NDUFS2, MRPL14) out of the 25 queried genes were not significantly associated with AD diagnosis or Braak stage, suggested broader biological impact of the Tau-seed interactors. Although we observed rescued eye phenotype and reduced seeding activities upon RNAi knockdown in *Drosophila*, the mRNA and protein levels of many of the hits are lower in AD brains and negatively associates with Braak stage, highlighting the complexity of Tau-seed interactors and suggesting factors involved in interactor levels or solubility could contribute to AD-related phenotypes.

**Figure 5.**
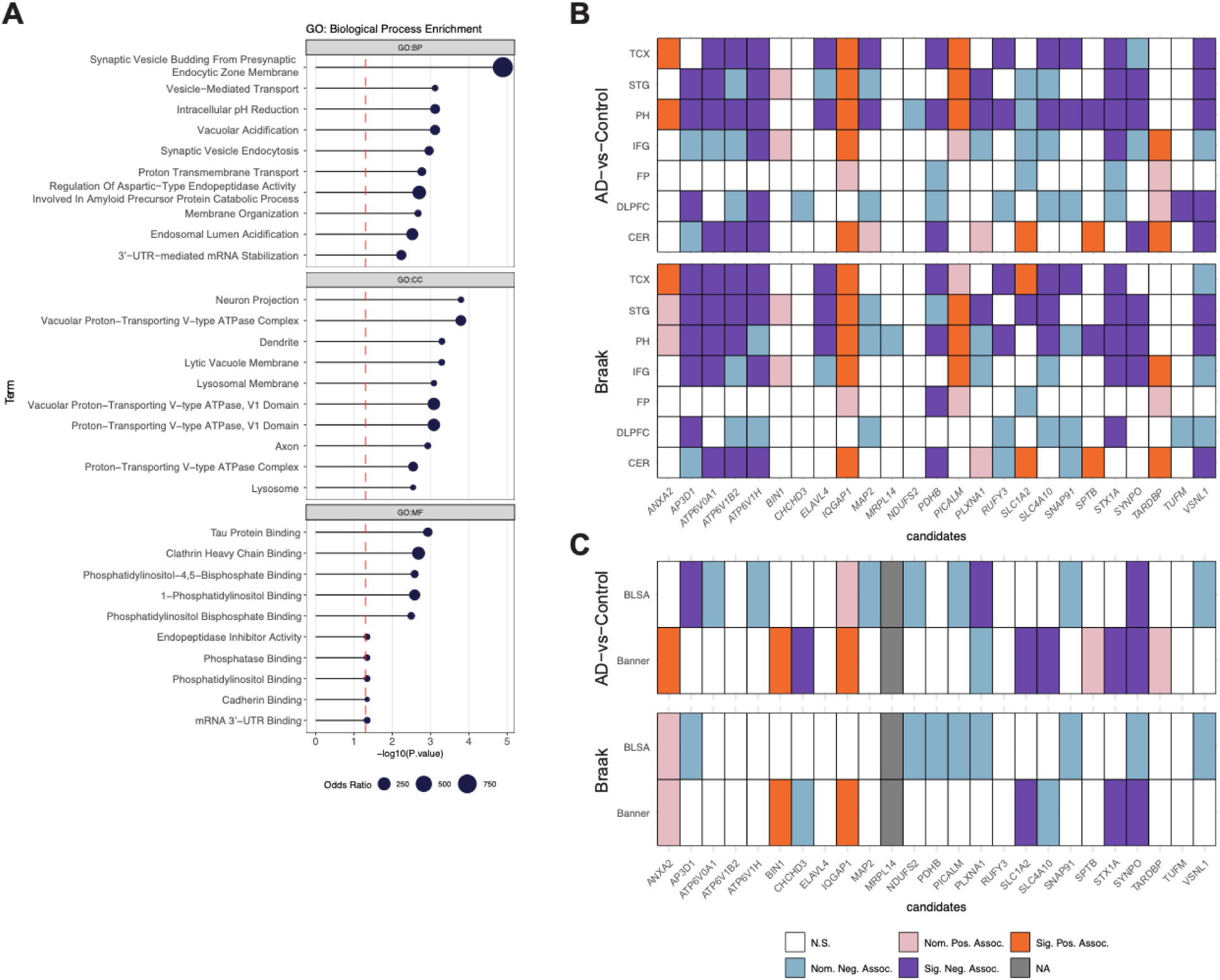
Top Tau-seed hits show consistent changes in post-mortem AD multi-omics datasets (A) Top 10 GO terms enriched among 25 Tau-seed interactors that reduced seeding and eye degeneration in *Drosophila*. (B) The association between RNA levels with AD diagnosis and Braak stage across seven RNA-seq datasets. (C) Protein abundance associations with AD diagnosis and Braak stage in Banner and BLSA proteomic datasets. Significance was defined as FDR-adjusted p value of less than 0.05, while nominal significance was defined as an unadjusted p-value<0.05.

### Functional analysis of key AD Tau-seed interactors in a tauopathy mouse model

To investigate the role of these candidates in the formation of Tau species with seeding activity in a mammalian system, we selected 11 interactor genes from the *Drosophila* screen for validation in the PS19 mouse model. Selection was guided by multiple criteria, with primary emphasis on eye degeneration and Tau-seeding activity observed in the *Drosophila* screen. Additional selection criteria included reported AD relevance in the literature, WGCNA module membership, known cellular functions related to synaptic, mitochondrial, or vesicle-associated pathways, and multi-omic correlation with at least one human AD dataset (Table S4). The selected Tau-seed interactors included TDP-43 (M3) and BIN1 (M2) due to their established AD relevance^35–37^, SNAP91 (M2), VSNL1 (M2), SYNPO (M3), and STX1A (M1) as synaptic-related proteins, ATP6V1B2 (M4), ATP6V1H (M3), and PICALM (M1) as vesicle-associated factors, and TUFM (M1) and NDUFS2 (M1) as mitochondria-related proteins. It is worthy to mention that while we prioritized eleven proteins for *in vivo* validation, the remaining candidates - part of a broader set of 25 modifiers-may also contribute to Tau toxicity and require future investigation. To downregulate gene interactors, neonatal PS19^38^ littermates were injected with AAV9 vectors carrying either a scrambled or an shRNA sequence targeting the mouse homolog of each of the eleven selected candidates. Four months post-injection, animals were euthanized, and soluble brain lysates were analyzed to assess Tau-seeding activity (Figure 6A). We observed a significant reduction in seeding activity upon downregulation of *Atp6v1h*, *Vsnl1*, *Stx1a*, *Bin1*, *Tufm*, *Picalm*, *Atp6v1b2*, and *Snap91* (Figure 6B). Surprisingly, downregulation of *Tardpb* resulted in a notable increase in seeding activity, suggesting a model-specific, differential role in Tau-seed formation, compared to *Drosophila* results. To assess the impact of the knock-down of Tau-seed interactors on total human Tau levels, we performed an MSD assay in PS19 brain lysates. Downregulation of *Stx1a*, *Picalm*, *Atp6v1b2*, and *Snap91* significantly reduced total human Tau levels, while other interactors had no significant effect on total Tau levels despite influencing Tau seeding (Figure 6C). Given the pathological relevance of pT231 and its strong correlation with AD HMW-Tau-seeding activity (Figure S3D), we next evaluated pT231 levels in the PS19 mice brains. Notably, *Stx1a*, *Bin1*, *Picalm*, *Atp6v1b2*, and *Snap91* knockdown significantly reduced pT231 levels (Figure 6D). To determine whether the interactors that have an effect on Tau-seeding activity influence HMW-Tau-seed formation, we fractionated shRNA-treated PS19 lysates by SEC and quantified seeding activity. As expected, knockdown of *Atp6v1h*, *Vsnl1*, *Stx1a*, *Bin1*, *Tufm*, *Picalm*, *Atp6v1b2*, and *Snap91* significantly reduced seeding activity in SEC fractions (Figure 6E). Interestingly, *Tardbp* knockdown caused a shift in seeding activity, evidenced by shared seeding between fractions F8 and F9, with a significant increase in F8 (Figure S5A, S5B), suggesting a potential role for TDP-43 in HMW-Tau-seed complex size. To evaluate how downregulation of Tau-seed interactors affects the formation and distribution of HMW-Tau, we analyzed total Tau levels in the HMW SEC fractions (F8, F9, F10). We observed significant reductions in these fractions in *Snap91*, *Stx1a*, *Picalm*, *Bin1*, *Tufm*, and *Atp6v1b2* knockdown mice, indicating that these proteins contribute to the formation of HMW-Tau complexes (Figure 6F). On the other side, *Atp6v1h* and *Vsnl1* downregulation do not influence the levels of total Tau in F9, suggesting that these interactors may affect the properties of Tau-seed by enhancing its seeding activity as previously suggested for BSN^11^. pT231 analysis in the same fractions revealed significant reductions of pT231 in *Snap91*, *Atp6v1h*, *Stx1a*, *Picalm*, *Bin1*, and *Tufm* knockdown mice (Figure 6G). No significant differences in pT231 levels were observed in *Atp6v1b2*, *Vsnl1* and *Tardbp*. Together, these findings highlight the heterogenous roles of Tau-seed interactors in regulating both HMW-Tau-seed formation, stabilization and modifications, providing new mechanistic insights into Tau pathology modulation.

**Figure 6.**
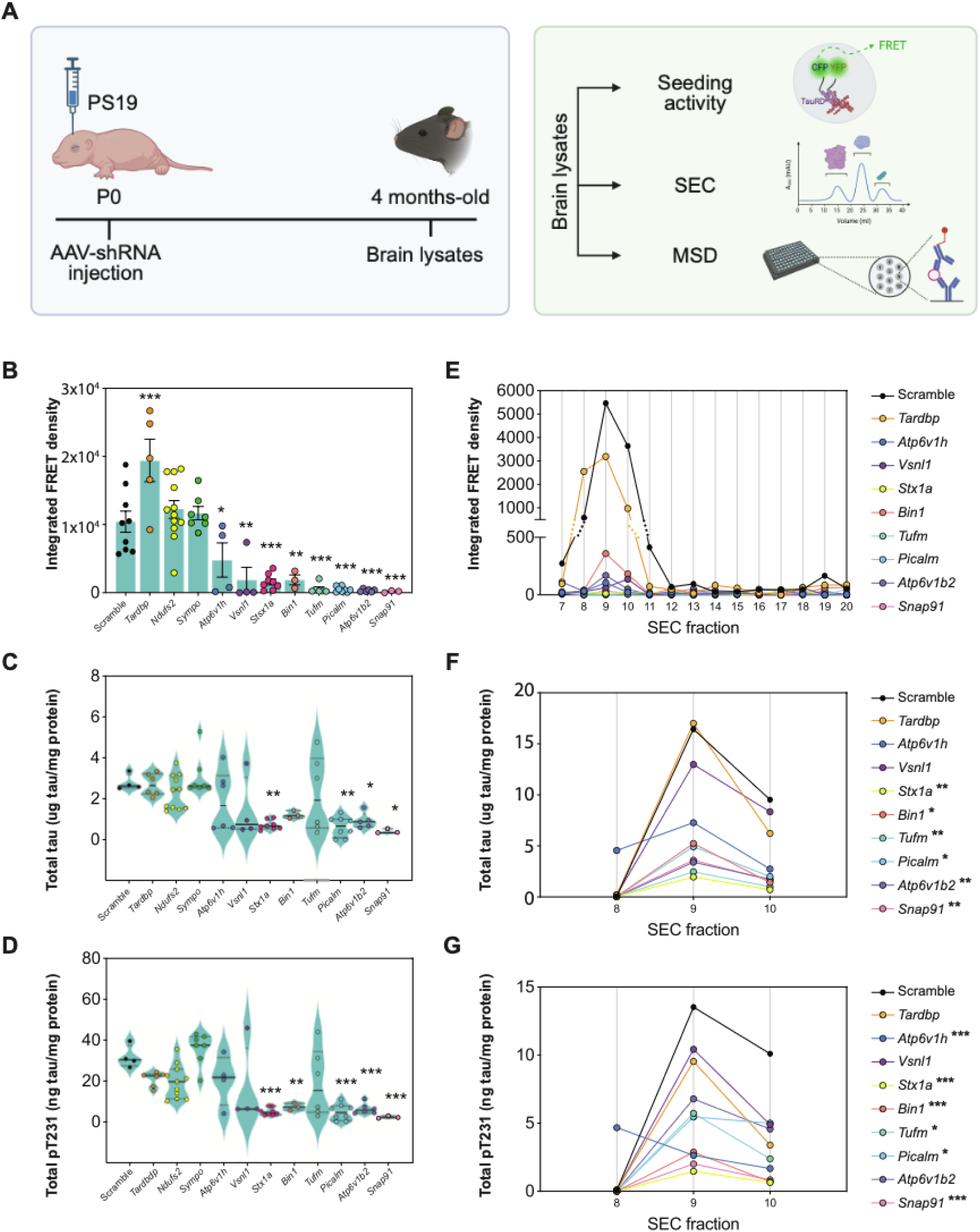
*In vivo* knockdown of Tau-seed interactors reduces pathological Tau in PS19 mice (A) Workflow for AAV-shRNA delivery targeting eleven Tau-seed interactors in PS19 mice at P0, followed by biochemical analysis at 4 months. Created in BioRender. (B-D) Quantification of Tau seeding activity (B), total Tau (C), and pT231 (D) by MSD in whole brain lysates. Data are shown as the mean ± s.e.m., and significance was determined by one-way ANOVA (*p<0.05, **p<0.005, ***p<0.0005; *n*=3-12). (E) Tau seeding activity in SEC fractions from PS19 brain lysates after shRNA treatment. Data are shown as the mean (*n*=3). (F and G) Total Tau (F) and pT231 (G) levels in SEC fractions 8, 9 and 10, measured by MSD following shRNA-mediated knockdown. Data are shown as the mean. Significance was determined by one-way ANOVA (*p<0.05, **p<0.005, ***p<0.0005; (*n*=3).

### HMW AD Tau-seed interactors exhibit distinct pathological co-deposition with Tau pathology

To investigate the pathological distribution of the key Tau-seed interactors in relation to pT231 Tau species, we performed immunofluorescence (IF) staining for pT231 in the middle frontal gyrus (MFG) of postmortem AD brains, along with Tau-seed interactors that either increased (TDP-43) or decreased (ATP6V1H, VSNL1, STX1A, BIN1, TUFM, PICALM, ATP6V1B2, and SNAP91) seeding activity in PS19 mice. Strikingly, each interactor exhibited a unique co-deposition pattern with pT231. While some interactors showed smaller, diffuse, or punctate accumulation (TDP-43, ATP6V1B2, SNAP91, VSNL1) others formed aggregated neuropil-like structures (ATP6V1H, BIN1, PICALM, STX1A, TUFM), suggesting an heterogenous association with Tau pathology (Figure 7). This variability could imply the existence of distinct Tau-seed complexes within the same brain that may contribute to varying pathogenic mechanisms, influencing disease progression and toxicity. The observed heterogeneity shows the complexity of Tau pathology in AD and raises the possibility that different Tau-seed populations may underlie specific disease trajectories.

**Figure 7.**
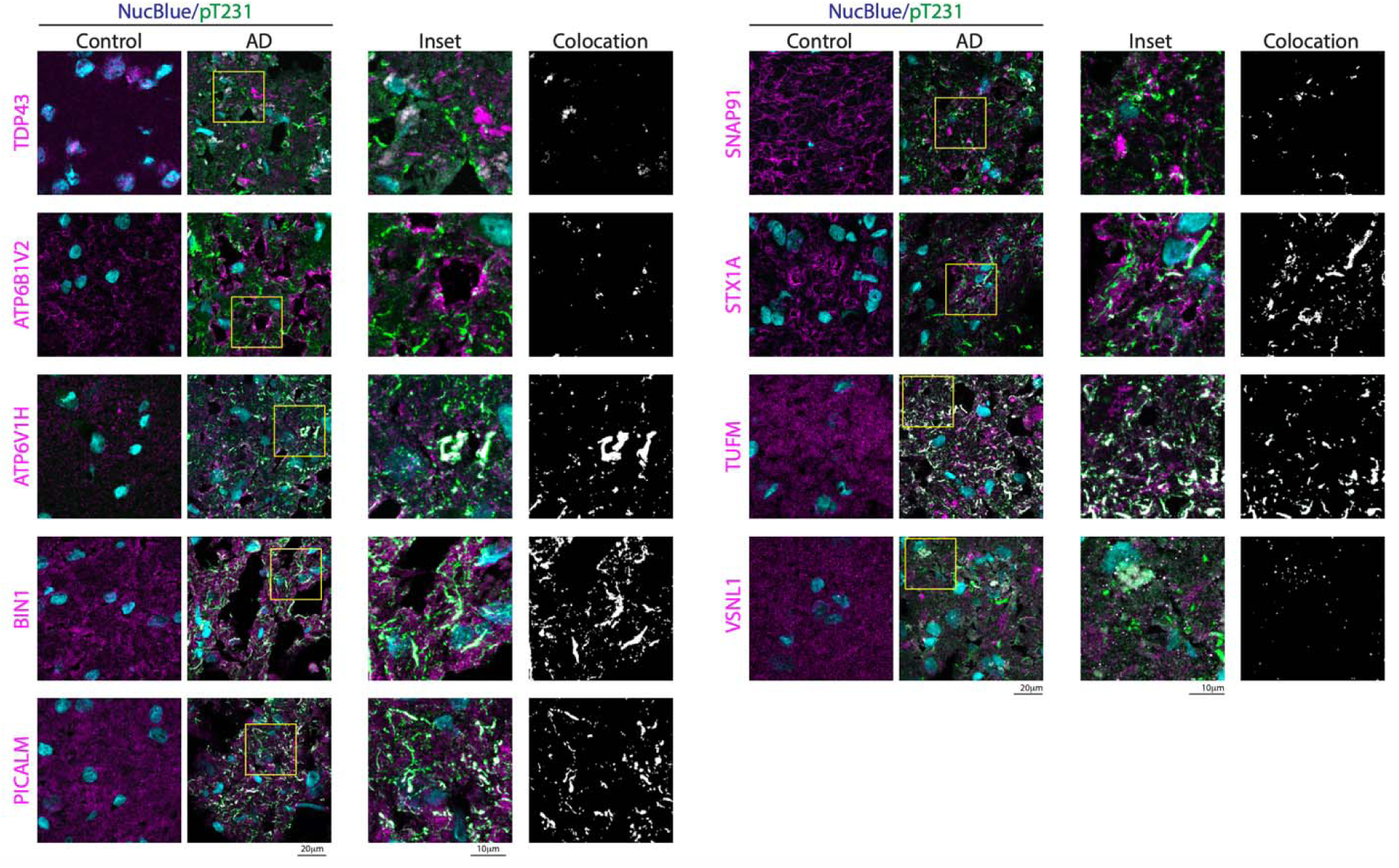
AD Tau-seed interactors colocalize with pathological pT231 in human brains (A) Representative immunofluorescence images and colocalization analysis in middle frontal gyrus brain sections from AD and age-matched control cases. Staining includes NucBlue (nuclei), pT231 (green), and Tau-seed interactors (magenta). Scale bars: 20 µm (main); 10 µm (insets).

## DISCUSSION

The understanding of pathological Tau aggregation and propagation mechanisms in the form of Tau-seed is a cornerstone to developing new strategies to combat neurodegenerative diseases such as AD and other Tauopathies. Here, we developed a systematic approach from proteomic to the use of different model systems to dissect the human Tau-seed interactome from AD and HMW-Tau from control brains. Our study shows that an HMW-Tau species is a driver of seeding activity in AD pathology, confirming previous results from our laboratory in a mouse model of tauopathy^11^. This AD interactome is characterized by distinct proteins and functional implications compared to non-seeding HMW-Tau derived from control cases. While previous studies have also identified a hyperphosphorylated HMW-Tau as the entity responsible for pathological propagation throughout the brain^9,10^, our findings extend this understanding by demonstrating that these aggregates are not composed solely of Tau. Instead, we have shown that this HMW-Tau requires specific interacting proteins to trigger the seeding capability, highlighting the role of Tau-associated proteins in disease pathogenesis. Notably, although control brains contain a comparable amount of Tau in this HMW fraction, they lack seeding activity, suggesting that qualitative differences in Tau aggregation and its associated interactome, rather than quantity alone, drive pathological seeding.

MS-based interactome profiling revealed a divergence between AD Tau-seed and control HMW-Tau interactors. While both share common cellular components and biological pathways, the AD Tau-seed interactome is uniquely enriched in axonal proteins. This finding aligns with the notion that axons serve as sites for Tau aggregation and propagation in AD^39,40^. Similar to other studies indicating that Tau binds synaptic vesicle proteins^16,41,42^, we observed an enrichment of synaptic vesicle proteins that bind Tau-seed. Furthermore, the presence of mitochondria- and synapse-related proteins suggests that Tau-seeds actively contribute to neurodegenerative cascades beyond passive aggregation, potentially exacerbating cellular dysfunction.

A comparative analysis of the AD and control interactomes highlights key functional distinctions. While control HMW-Tau interactors appear to form relatively homogeneous and weakly correlated networks, visual inspection of the eigengene heatmap and the correlation network of co-expression modules suggests that AD Tau-seed interactors are organized into more densely connected modules with higher internal correlation, enriched in mitochondrial function, synaptic vesicle dynamics, and intracellular vesicle-mediated trafficking. Notably, module correlations also revealed the formation of two distinct, module clusters that are negatively correlated to each other. One cluster linked mitochondrial and synaptic vesicle dynamics, while the other associated intracellular trafficking with ribosomal function and macromolecular disassembly. This modular organization could suggest the presence of at least two Tau-seed strains, each comprising distinct sets of interactors from diverse subcellular localizations and compartments. These findings align with the emerging hypothesis proposing that Tau-seed populations are heterogeneous across individuals and that their conformational and pathological properties may be shaped by specific cellular contexts and interacting proteins^43–46^. This restructuring suggests that Tau-seeds are not merely products of neurodegeneration but active participants in modifying key cellular processes, potentially amplifying disease pathology.

Using a *Drosophila* tauopathy model, we functionally validated AD Tau-seed interactors and identified 25 proteins that reduced Tau toxicity and seeding activity. Network analysis categorized these interactors into three primary clusters: synaptic, mitochondrial, and vesicle-related processes. Multi-omic integration with human AD datasets further supports the pathological relevance of the identified Tau-seed interactors, which are consistently dysregulated at both the RNA and protein levels across multiple brain regions. Notably, eight protein interactors (ATP6V1H, VSNL1, STX1A, BIN1, TUFM, PICALM, ATP6V1B2, and SNAP91) reduced Tau seeding activity after knockdown in PS19 mice. Several of these proteins showed significant correlation with AD diagnosis and Braak stage across transcriptomic and proteomic datasets. While we prioritized 11 proteins for *in vivo* validation in PS19 mice based on the *Drosophila* screen and relevance to human AD datasets, this selection does not preclude the potential relevance of the remaining candidates. The broader set of 25 proteins that reduced pathology in flies warrants further investigation, and future studies should evaluate additional mechanisms and candidate interactors that may contribute to Tau toxicity in distinct biological contexts. Interestingly, while all eight validated interactors reduced Tau seeding activity in PS19 mice, their effects on total Tau and pT231 levels were not uniform. STX1A, TUFM, ATP6V1B2, and SNAP91 significantly reduced both total Tau levels and pT231 as measured by MSD, whereas BIN1reduced only pT231. These differences suggest that reductions in seeding activity are not necessarily coupled with decreased overall Tau or specific pathological phosphorylations. This divergence indicates that these interactors may influence distinct steps in Tau aggregation, post-translational modification, or the assembly of HMW-Tau species with seeding activity. Together, these findings support a model in which seeding activity, total Tau abundance, and pathological phospho-epitopes are modulated by partially overlapping but mechanistically distinct pathways.

These mechanistic differences likely reflect the diverse cellular pathways through which these interactors influence Tau pathology. In particular, STX1A and SNAP91 are core components of the presynaptic vesicle fusion machinery^47,48^. Their relevance is supported by accumulating evidence that secreted Tau can propagate among synaptically connected neurons^49,50^. Furthermore, recent work by Tracy et al. demonstrated that wild-type Tau interacts with numerous presynaptic proteins at the active zone in human iPSC-derived neurons^6^. This interaction was enhanced by neuronal activity, suggesting that synaptic vesicle release machinery directly participates in activity-dependent Tau secretion through protein-protein interactions. Thus, the downregulation of STX1A and SNAP91 may reduce presynaptic Tau-seed release, limiting extracellular seed availability and trans-synaptic propagation. Other validated candidates, such as ATP6V1H and ATP6V1B2, encode subunits of the vacuolar ATPase (v-ATPase), which is essential for lysosomal acidification and autophagic flux^51^. Although impairment of v-ATPase can compromise Tau degradation^52,53^, partial suppression may attenuate endosomal trafficking or fusion steps required for Tau-seed release, thereby reducing seeding potential. Similarly, VSNL1, a calcium-binding neuronal protein, is a known cerebrospinal fluid biomarker in AD^54^ and is involved in calcium-mediated synaptic signaling; its knockdown may attenuate calcium-mediated synaptic activity and reduce Tau-seed secretion, consistent with previous findings implicating neuronal activity in Tau release^49^. Furthermore, TUFM, a mitochondrial elongation factor involved in mitochondrial translation and proteostasis^55^, is also reduced in AD brains. It may influence Tau pathology by modulating cellular stress responses and protein folding quality control. Together, these data suggest that presynaptic machinery, vesicular trafficking, and mitochondrial pathways contribute to the regulation of Tau seeding activity and that perturbation of these systems can mitigate the spread of pathological Tau *in vivo*.

On the other hand, PICALM and BIN1, both established genetic risk factors for AD^56^, were positively correlated with disease in transcriptomic and proteomic AD datasets. These proteins are key regulators of endocytosis and membrane trafficking^57,58^.

Mechanistically, PICALM facilitates Tau uptake and clearance^59^, whereas BIN1 has been shown to influence Tau pathology by modulating endocytic flux^60^. *In vitro* studies have shown that BIN1 knockdown promotes Tau propagation via enhanced endocytosis and trafficking^60^. However, our *in vivo* screening in *Drosophila* and PS19 mice show that BIN1 downregulation reduces Tau seeding activity and phosphorylation at pT231. These findings align with recent *in vivo* work where conditional BIN1 loss reduced Tau pathology in the hippocampus^35^. This discrepancy with *in vitro* studies likely reflects model-specific differences in cellular complexity and regional context. BIN1’s role is isoform- and cell-type-dependent, particularly between neurons and oligodendrocytes^61,62^.

Conversely to the *Drosophila* screening, TDP-43 downregulation paradoxically increased seeding in PS19 mice, in a model-specific interaction manner, suggesting a potential protective function in mice, as supported by prior studies on TDP-43 knockdown^37^. These findings underscore the diverse contributions of Tau interactors to disease progression and suggest that selectively targeting specific interactors could have differential effects on Tau pathology. Importantly, our results also raise the possibility that a combinatorial or multi-targeting approach-rather than focusing on a single interactor-may offer enhanced therapeutic benefits. Given that several interactors converge on interconnected pathways, simultaneously modulating multiple nodes within the Tau-seed network could more effectively suppress seeding activity and slow disease progression.

The differential co-localization patterns of Tau-seed interactors with pT231 in postmortem AD brains could suggest that Tau pathology is mediated through diverse molecular assemblies. The presence of both diffuse puncta and neuropil-like aggregates implies that distinct interactors may contribute to pathogenic processes via separable mechanisms, potentially reflecting region or cell type-specific vulnerabilities. Given the strong correlation between pT231 and human seeding activity, these findings raise the possibility that specific HMW-Tau-seed complexes differentially modulate disease pathology. This heterogeneity underscores the multifactorial nature of Tau aggregation and highlights the need to re-examine the pathways through which Tau toxicity is initiated and propagated.

Our study provides a detailed resource of Tau-seed–specific protein interactions and their functional consequences, offering mechanistic insight into how pathological Tau species, which represent less than 5% of total Tau, engage with cellular pathways in the human brain. By integrating proteomics, functional genetics, and *in vivo* validation in two model system, we delineate molecular networks that may modulate Tau seeding and propagation. The distinct interactomes, network reorganization, and functional consequences of Tau-seed interactors suggest that Tau pathology extends beyond fibrillar aggregation, including a broader landscape of protein interactions and cellular dysfunctions. These findings establish a basis for targeted strategies aimed at disrupting Tau-seed formation or stabilization to attenuate neurodegenerative progression in AD.

## Data and code availability

Mass spectrometry data have been deposited in the ProteomeXchange Consortium member MassIVE with accession MSV000095940. Data are currently password protected and can be accessed at https://massive.ucsd.edu/. Source data are provided with this paper. All other numerical data are available from the corresponding author upon reasonable request. Human post-mortem transcriptomic and proteomic data can be retrieved following the accession number listed in the Key Resource Table section.

## ACKNOWLEDGMENTS

This work was supported by NINDS 1R01NS119280, NIA 1RF1AG059639 grants, and ALZDISCOVERY-1049108 from the Alzheimer’s Association grant; the Cure Alzheimer’s Foundation and the Rainwater Charitable Foundation to C.A.L.-R., AARFD-21-847663 to P.M., NIA P30AG066507 (Johns Hopkins ADRC) to J.T, and 1R01AG080917-01 and 1R21NS135424, as well as a CTSI-PDT grant, to J.-C.R This publication was also supported by the Sarah Roush Memorial Fellowship in Alzheimer’s Disease, the Indiana Alzheimer’s Disease Research Center, and the Stark Neurosciences Research Institute and made possible by the Indiana Clinical and Translational Sciences Institute, funded in part by the grant # UL1TR002529 to P.M. from the National Institutes of Health, National Center for Advancing Translational Sciences. This work was supported by National Institute on Aging (RF1 AG051504, U01 AG046139 R01 AG061796, and U19 AG074879 to N.E.T). N.E.T. is also supported by the Alzheimer’s Association Zenith Fellows Award (ZEN-22-969810).

The mass spectrometry work performed in this work was done by the Indiana University School of Medicine Proteomics Core. Acquisition of the IUSM Proteomics core instrumentation used for this project was provided by the Indiana University Precision Health Initiative. The proteomics work was supported, in part, by the Indiana Clinical and Translational Sciences Institute (funded in part by Award Number UL1TR002529 from the National Institutes of Health, National Center for Advancing Translational Sciences, Clinical and Translational Sciences Award) and, in part, by the IU Simon Comprehensive Cancer Center Support Grant (Award Number P30CA082709 from the National Cancer Institute).

AD Knowledge Portal: The results published here are in whole or in part based on data obtained from the AD Knowledge Portal (https://adknowledgeportal.org). AMP-AD datasets: The results published here are in whole or in part based on data obtained from the AMP-AD Knowledge Portal (https://doi.org/10.7303/syn2580853). Mayo Clinic: The Mayo RNAseq study data was led by Dr. Nilüfer Ertekin-Taner, Mayo Clinic, Jacksonville, FL as part of the multi-PI U01 AG046139 (MPIs Golde, Ertekin-Taner, Younkin, Price). Samples were provided from the following sources: The Mayo Clinic Brain Bank and Banner Sun Health Research Institute. Data collection was supported through funding by NIA grants P50 AG016574, R01 AG032990, U01 AG046139, R01 AG018023, U01 AG006576, U01 AG006786, R01 AG025711, R01 AG017216, R01 AG003949, NINDS grant R01 NS080820, CurePSP Foundation, and support from Mayo Foundation. Study data includes samples collected through the Sun Health Research Institute Brain and Body Donation Program of Sun City, Arizona. The Brain and Body Donation Program is supported by the National Institute of Neurological Disorders and Stroke (U24 NS072026 National Brain and Tissue Resource for Parkinsons Disease and Related Disorders), the National Institute on Aging (P30 AG19610 Arizona Alzheimer’s Disease Core Center), the Arizona Department of Health Services (contract 211002, Arizona Alzheimer’s Research Center), the Arizona Biomedical Research Commission (contracts 4001, 0011, 05-901 and 1001 to the Arizona Parkinson’s Disease Consortium) and the Michael J. Fox Foundation for Parkinsons Research. MSBB: These data were generated from postmortem brain tissue collected through the Mount Sinai VA Medical Center Brain Bank and were provided by Dr. Eric Schadt from Mount Sinai School of Medicine. ROSMAP: Study data were provided by the Rush Alzheimer’s Disease Center, Rush University Medical Center, Chicago. Data collection was supported through funding by NIA grants P30AG10161 (ROS), R01AG15819 (ROSMAP; genomics and RNAseq), R01AG17917 (MAP), R01AG30146, R01AG36042 (5hC methylation, ATACseq), RC2AG036547 (H3K9Ac), R01AG36836 (RNAseq), R01AG48015 (monocyte RNAseq) RF1AG57473 (single nucleus RNAseq), U01AG32984 (genomic and whole exome sequencing), U01AG46152 (ROSMAP AMP-AD, targeted proteomics), U01AG46161(TMT proteomics), U01AG61356 (whole genome sequencing, targeted proteomics, ROSMAP AMP-AD), the Illinois Department of Public Health (ROSMAP), and the Translational Genomics Research Institute (genomic).

## AUTHOR CONTRIBUTIONS

P.M. and C.A.L.-R. conceived and designed the project with the contribution of H.P., Y.Y., D.L., A.A., N.J-G., Y.M., N.E.-T. and S.D., performing experiments. J.R.-O. and J.T. characterized and provided AD and control cases. S.D. and J-C.R. performed EM analysis. A.A., Y.M. and N.E.-T. conducted bioinformatic analysis. P.M. and D.L. conducted *Drosophila* experiments. P.M., H.P., Y.Y. and A.A. conducted screening experiments in mice. P.M. and N.J.-G. performed AD cases immunofluorescence experiments and analysis. P.M. wrote the manuscript with contributions from all the authors and supervision of C.A.L.-R.

## MATERIALS AND METHODS

### Cell Culture

For seeding activity assays, we used the TauRD P301S FRET Biosensor cells, commercially available (American Type Culture Collection, ATCC).

### Mice

All experiments involving mice followed the guidelines outlined in the Guide for the Care and Use of Laboratory Animals by the National Institutes of Health (NIH) and received approval from the Institutional Animal Care and Use Committee at the Indiana University School of Medicine (IUSM). Mice were bred and housed at the IUSM animal care facility, adhering to US Department of Agriculture standards, which included a 12-hour light/dark cycle, access to food and water *ad libitum*, a temperature of 25°C, and humidity between 40–60%, in accordance with the NIH guidelines. The PS19 mouse model, which overexpresses human 1N4R Tau with the P301S mutation on a C57B6/J background, was acquired directly from The Jackson Laboratory. In all experiments, 4-month-old and their wild-type (WT) littermates of both sexes, were used. Mice were randomly assigned to treatment and experimental groups.

### Drosophila

*D. melanogaster* stocks and genetic crosses were maintained on Fly Food B (LabExpress) at 25°C under a 12-hour light/dark cycle. All fly experiments were conducted 10 days post-eclosion. All transgenic flies were obtained from Bloomington *Drosophila* Stock Center (BDSC).

### Human Subjects

Frozen samples consisting of blocks of postmortem brain tissues from individuals with AD and age-matched healthy controls were provided by the Brain Resource Center at Johns Hopkins. AD cases had severe AD neuropathological change, and Braak neurofibrillary stage V–VI. See details in Figure S1A.

### Tris-buffered saline-soluble homogenates

Each brain tissue sample was homogenized in TBS buffer (Corning) at a 1:10 (weight/volume) ratio, with the addition of a protease inhibitor cocktail (Thermo Scientific). The samples were centrifuged at maximum speed (14,000 rpm) for 15 minutes at 4°C. The resulting supernatants were divided into aliquots, snap-frozen, and stored at −80°C until further analysis.

### Size exclusion chromatography

Size-exclusion chromatography (SEC) was carried out using a Superose 6 Increase 10/300 GL column (GE Healthcare, 29091596) on an ÄKTA pure 25 L chromatography system (GE Healthcare, 29018224). The column was equilibrated with 1.5 column volumes of a buffer containing 50 mM NaCl and 50 mM Tris at pH 8.0, with a flow rate of 0.7 ml/min. Samples were clarified by centrifugation at 10,000g for 10 minutes. Protein concentration was measured using a Bradford assay (Thermo Scientific), and depending on the sample, 1–5 mg of total protein from the supernatant was used for separation. The supernatant was concentrated to approximately 200 μl using a 0.5-ml 3K Amicon centrifugal filter (Millipore Sigma) before being loaded onto the column through sample loop injection. Starting at the point of injection, 1-ml fractions were collected at a flow rate of 0.3 ml/min into tubes containing EDTA-free protease inhibitor (Thermo Scientific).

### Tau-seeding assay

The seeding assay was conducted following our previously described protocol^8,22^. In brief, TauRD P301S FRET Biosensor cells (ATCC) were seeded at 35,000 cells per well in 130 μl of medium in a 96-well plate and incubated overnight at 37°C. The next day, cells were transfected with 20 μg of total protein from cell or brain lysate/10µl SEC fraction per well using Lipofectamine 2000 and incubated for an additional 48 hours at 37°C. Cells were then collected by trypsinization, and flow cytometry was performed using a BD LSRFortessa X-20 equipped with a High Throughput Sampler. Data acquisition utilized BD FACS Diva (v8.0) software, and analysis was carried out with FlowJo (v10.0). The BV421 channel (405 nm excitation, 450/50 nm emission) was used to detect CFP, while the BV510 channel (405 nm excitation, 525/50 nm emission + 505LP) was employed to measure the FRET signal, with compensation applied to minimize CFP spillover into the FRET channel. FlowJo was used for data analysis, following the gating strategy outlined in Figure S1B. Seeding was quantified by integrated FRET density, which is calculated as the product of the percentage of FRET-positive cells and the median fluorescence intensity of those FRET-positive cells.

### Human Tau ELISA

ELISA was conducted on SEC fractions using the Tau (Total) Human ELISA Kit (Invitrogen) according to the manufacturer’s instructions. Lysates were diluted 1:50,000 in blocking buffer. Fractions F7–F14 were diluted at a 1:2,000 ratio in blocking buffer, while fractions F15–F22 were diluted at a 1:20,000 ratio in blocking buffer.

### Immunoprecipitation

Tau was immunoprecipitated from AD and control brain samples, using 2 μg of biotinylated HT7 antibody (Thermo Fisher) for every 100 ng of human Tau, as quantified by ELISA. For comparison, IgG isotype control antibodies (BioLegend) were used. Both the immunoprecipitated (IP) and flow-through samples were then subjected to further analysis.

### Electron microscopy

Immunoprecipitated samples were examined using negative-stain biological transmission electron microscopy (EM), following a previously described protocol^67^. In brief, 3 μl of the IP sample was directly applied to discharged carbon-coated copper TEM grids and allowed to incubate for 1 minute. The grids were gently rinsed with deionized water, ensuring they did not dry out, and stained with 3.5 μl of 1% (wt/vol) phosphotungstic acid solution for 1 minute. Excess solution was blotted off using Whatman filter paper. Imaging was performed with an FEI Tecnai T12 TEM operating at 80 kV, and images were captured using Gatan Digital Micrograph software. Measurements of width were conducted using ImageJ (NIH). Three independent samples were analyzed, and the results were plotted using Prism 9.0 software.

### Mass spectrometry

Sample preparation, mass spectrometry analysis, bioinformatics, and data evaluation for quantitative proteomics and phospho-proteomics experiments were performed in collaboration with the Indiana University Proteomics Center for Proteome Analysis at the Indiana University School of Medicine similarly to previously published protocols^68^.

### Sample Preparation

On bead samples were submitted to the IUSM Center for proteome analysis where proteins were denatured in 8 M urea, 100 mM Tris-HCl, pH 8.5 and reduced with 5 mM tris(2-carboxyethyl)phosphine hydrochloride (TCEP, Sigma-Aldrich) for 30 minutes at room temperature. Samples were then alkylated with 10 mM chloroacetamide (CAA, Sigma Aldrich) for 30 min at room temperature in the dark, prior to dilution with 50 mM Tris. HCl, pH 8.5 to a final urea concentration of 2 M for Trypsin/Lys-C based overnight protein digestion at 37 °C (0.5 µg protease, Mass Spectrometry grade, Promega)

### Peptide Purification and Labeling

Digestions were acidified with trifluoroacetic acid (TFA, Sigma Aldrich, 0.5% v/v) and desalted on 50 mg waters Sep-Pak cartridges (Waters™) with a wash of 1 mL of 0.5% TFA followed by elution in 3x 200 µL 70% acetonitrile 0.1% formic acid (FA).

### Tandem Mass Tag labeling

Dried peptides were resuspended in 25 µL of 100 mM Triethylammonium bicarbonate, pH 8.5 (diluted from Thermo Scientific™) and labeled with 0.25 mg of TMTpro reagent for 2 hours at room temperature (see manufacturer’s instructions Thermo Scientific). Reactions were quenched with 0.3 % final v/v hydroxylamine, and samples from each antibody group were pooled and dried by speed vacuum. The 4 grouped samples were each resuspended in 0.5% TFA and excess TMTpro reagent was removed using a Waters Sep-Pak Vac cartridge (Waters) with a 1 mL wash of water followed by 1 mL wash of 5% acetonitrile 0.1% FA and elution in 70% ACN 0.1% FA. Elutions were dried and resuspended in 25 µL 0.1% FA.

### Nano-LC-MS/MS

Mass spectrometry was performed utilizing an EASY-nLC 1200 HPLC system (Thermo Scientific) coupled to Eclipse orbitrap™ mass spectrometer with FAIMSpro interface (Thermo Scientific). 1/5 of each fraction was loaded onto a 25 cm Aurora Ultimate column (Ion Opticks) at 350 nL/min. The gradient was increased from 5-35% B over 160 minutes; 35-95% B over 10 mins; held at 95% for 2 minutes; and dropping from 80-5% B over the final 5 min (Mobile phases A: 0.1% FA, water; B: 0.1% FA, 95% Acetonitrile (Fisher Scientific). The mass spectrometer was operated in positive ion mode, default charge state of 2, advanced peak determination on, and EASY-IC. Three FAIMS CVs were utilized (−45 CV; −55 CV; −65 CV) each with a cycle time of 1 s and with identical MS and MS2 parameters. Precursor scans (m/z 400-1600) were done with an orbitrap resolution of 120000, RF lens% 30, 50 ms maximum inject time, standard AGC target, minimum MS2 intensity threshold of 2.5e4, MIPS mode set to peptide, including charges of 2 to 6 for fragmentation with 60 sec shared dynamic exclusion. MS2 scans were performed with a quadrupole isolation window of 0.7 m/z, 34% HCD collision energy, 50000 resolution, 200% AGC target, dynamic maximum IT, fixed first mass of 100 m/z.

### Mass spectrometry Data Analysis

Resulting RAW files were analyzed in Proteome Discover 2.5 (Thermo Fisher Scientific) with a *H. sapiens* reference proteome FASTA (downloaded from Uniprot 051322 with 78806 entries) plus common contaminants (73 entries^69^). SEQUEST HT searches were conducted with a maximum number of 3 missed cleavages; precursor mass tolerance of 10 ppm, and a fragment mass tolerance of 0.02 Da. Static modifications used for the search were carbamidomethylation on cysteine (C). Dynamic modifications included and oxidation of methionine (M), deamidation of asparagine or arginine, phosphorylation on serine, threonine or tyrosine, and acetylation, methionine loss, or methionine loss plus acetylation on protein N-termini. Percolator False Discovery Rate was set to a strict peptide spectral match FDR setting of 0.01 and a relaxed setting of 0.05. Results were loaded into Scaffold Q+S 5.2.2 (Proteome Software) for viewing. 1851 proteins were identified with at least 1 peptide.

### Interactor identification

For each biological replicate, abundance of HT7 IP was divided by abundance of IgG IP and result was named “IP enrichment ratio”. One-tailed one-sample t-test was performed on the hypothesis log2(IP enrichment ratio) > 0 across biological replicates to generate p-values. Samples that contained zero or missing abundances were omitted from the test. For each group (Ctrl and AD) separately, BKY 2-stage stepup was used to control FDR at the 5% level^70^. Proteins with q < 0.05 were taken as Tau interactors.

### WGCNA analysis and protein correlation network visualization

#### WGCNA

WGCNA analysis was performed according to previous literature, with modifications^27^. Starting from raw mass spec abundances, protein abundances were normalized to the sum of abundances for each sample. Proteins that have zero and N/A abundance values were omitted. WGCNA was performed on the log2 of HT7 IP abundances for each group (Ctrl and AD) separately, using the following parameters

- corType = “bicor”
- maxPOutliers = 0.05
- power = 10
- networkType = “signed hybrid”
- TOMType = “signed”
- minModuleSize = 10
- pamRespectsDendro = FALSE
- reassignThreshold = 0
- mergeCutHeight = 0.25

Then, for protein correlation network visualization, WGCNA module assignments and network adjacency values were imported into Cytoscape^66^. Nodes (proteins) were colored by module for pre-visualization. Lay out on nodes was performed using Edge-weighted Force directed (BioLayout) algorithm, using adjacencies as weights. Parameters were adjusted until modules are well clustered together, and the nodes do not look like a crystaline lattice but have a more amorphous structure. The following settings were used for Ctrl and AD networks:

- Edge column that contains the weights = adjacency
- How to interpret weight values = normalized value
- The minimum edge weight to consider = 0
- The maximum edge weight to consider = 1
- The default edge weight to consider = 0.5
- Divisor to calculate the attraction force = 10
- Multiplier to calculate the repulsion force = 5
- Multiplier to calculate the gravity force = 0
- Constant force applied to avoid conflicts = 0
- Percent of graph use for node repulsion calculations = 100
- Amount of extra room for layout = 10
- Initial temperature = 1e3
- Number of iterations = 1e5
- Don’t partition graph before layout = checked
- Randomize graph before layout = checked
- Layout nodes in 3D = unchecked

Final rendering was done externally since Cytoscape has no ability to specify drawing order of individual elements. The network was exported as .cx or .cx2 file to export node positions. Then an SVG file was generated using node positions from Cytoscape layout. Nodes were colored by module and sized by eigenvector centrality to the module it belongs. Graph edges were shaded using a perceptually linear colormap, opacity of each edge was set to the adjacency value.

The rendering z-order of edges are layered by increasing adjacency, with lowest adjacency edges on the bottom, and highest adjacency on the top. All nodes are rendered on top of all edges.

### AD vs control differential protein abundance

To identify Tau interactors that have a different abundance in AD and control brains, we compared the protein abundance using the proteomic data from the HT7 fraction.

Briefly, proteins that were detected only in AD or Control were first excluded. The LFQ data were then normalized to total protein abundance for each sample and log transformed. To account for the missing values, proteins that are detected in less than 50% of the samples within each diagnosis group is removed. Differential protein abundance was calculated using a two-sided t-test. The enriched Gene Ontology (GO) terms were queried using enrichR^65^. GO enrichment analysis were performed separately for interactors that are more abundant (Up in AD) or less abundant (Down in AD), using a nominally significant (unadjusted p-values < 0.05) cut off.

### External Datasets

Gene expression analysis was performed using published bulk brain RNAseq datasets generated from Mayo Clinic (Mayo)^28^, Mount Sinai School of Medicine (MSSM)^30^, and Religious Orders Study/Memory and Aging Project (ROSMAP)^29^ (Key Resources Table). Based on the distribution of the CQN-normalized gene expression, lowly expressed genes were excluded. The associations between CQN-normalized gene expression and either AD diagnosis or Braak stage was carried out using a linear mixed-effect model while adjusting for technical and biological covariates. We treated age at death, sex, RIN as fixed effect variables, while sequencing batch was treated as a random effect variable. The RNAseq data in ROSMAP and Mayo Clinic datasets were generated using brain tissue from different brain banks, which were also adjusted. Participants from the Mayo Clinic dataset received pathological diagnosis of AD or control based on Braak stage ≥4 or ≤3, respectively. Controls from Mayo Clinic Brain Bank lacked any neuropathological diagnosis of other neurodegenerative diseases. As previously published^71,72^. Mount Sinai and ROSMAP AD cases had Braak stage of ≥4. Mount Sinai AD donors had CERAD score of possible, probable or definite AD, and ROSMAP AD donors had CERAD of probable or definite AD. Controls from Mount Sinai and ROSMAP were defined as patients with Braak stage of less than or equal to III and having a CERAD score lower than defined for AD from respective cohorts. Donor samples that did not meet criteria for either AD or control were excluded from the AD-vs-Control analysis. For the association between gene expression and Braak stages, all available donors with Braak scores were included. Multiple testing was corrected for using the false discovery rate.

The Baltimore Longitudinal Study on Aging (BLSA)^32^ study and the Banner Sun Health Research Institute (Banner)^33^ study proteomics datasets were previously published and retrieved from AD Knowledge Portal (Key Resources Table). The LFQ intensity were log2 transformed and normalized to median intensity for each protein. Missing values were imputed within each sample based on a normal distribution with the mean of which is at the 10th percentile of the overall intensity, and the standard deviation is the 30th percentile of the standard deviation of the overall intensity. Peptides with over 50% imputed values were excluded from the analysis. The associations between peptide abundance and AD diagnosis or Braak stage were evaluated using a linear model while adjusting for covariates including sex, age at death, and post-mortem interval (PMI). AD or control diagnosis were defined based on the Braak and CERAD scores using the same criteria for the RNAseq datasets. Multiple testing was adjusted for using the false discovery rate methods on all peptides within a dataset.

### Drosophila Screening

Transgene overexpression was achieved using the Gal4/UAS system, utilizing the GMR-Gal4 and GMR-Tau^P301L^ lines obtained from the Bloomington *Drosophila* Stock Center. A total of 115 transgenic fly lines, used for screening the AD Tau-seed interactome, were sourced from the BDSC (Figure S4A and Key resource table). Crosses targeting Tau-seeds were performed by mating Tau^P301L^; GMR-Gal4 flies with UAS-RNAi flies. To evaluate eye phenotypes and assess seeding, adult flies were immobilized by freezing at −80°C and then prepared for light microscopy. Imaging was conducted with a Leica DMC6200 camera equipped with a 10x objective, using white light to illuminate each ommatidium until a single reflection spot was centered. Eye phenotype analysis was performed using the Flynotyper plug-in on ImageJ, according to established protocols^73^.

### Adeno-associated virus production and injections

For Tau-seed interactor candidates’ downregulation, neonatal (P0) PS19 and WT mice were injected with the AAV scramble or shRNA sequences. Animals were euthanized at 4 months after injections. shRNA sequences were cloned downstream of the U6 promoter and packaged into AAV9 from VectorBuilder.

### Immunofluorescence on human brain frozen sections

Frozen human brain sections were fixed in 4% paraformaldehyde at room temperature (RT) for 1 hour, followed by permeabilization in 0.01% Triton X-100 in TBS for 1 hour. Antigen retrieval was performed by heating sections at 95°C for 10 minutes in a high pH antigen retrieval solution using a microwave oven. Sections were washed twice with TBS (5 minutes each) and incubated with TrueBlack (Biotium) reagent for 3 minutes to reduce autofluorescence, followed by three PBS washes. Tissues were blocked in TBS containing 10% goat serum and 0.01% Triton X-100 for 1 hour at RT.

Serial immunofluorescence was performed as follows: On day 1, sections were incubated overnight at 4°C with the first primary antibody diluted in blocking solution. On day 2, sections were washed three times with TBS, then incubated with the corresponding secondary antibody diluted in blocking solution for 2.5 hours at RT. Following three washes in PBS, sections were blocked again, and the second primary antibody was applied overnight at 4°C. On day 3, sections were washed three times with TBS and incubated with the secondary antibody against the second primary antibody for 2.5 hours at RT. After three final PBS washes, sections were mounted with Fluoromount (Invitrogen).

## QUANTIFICATION AND STATISTICAL ANALYSIS

The statistical methods used for each experiment are detailed in the corresponding sections of the Materials and Methods and/or in the figure legends.

## Supplementary figures

**Figure S1.**
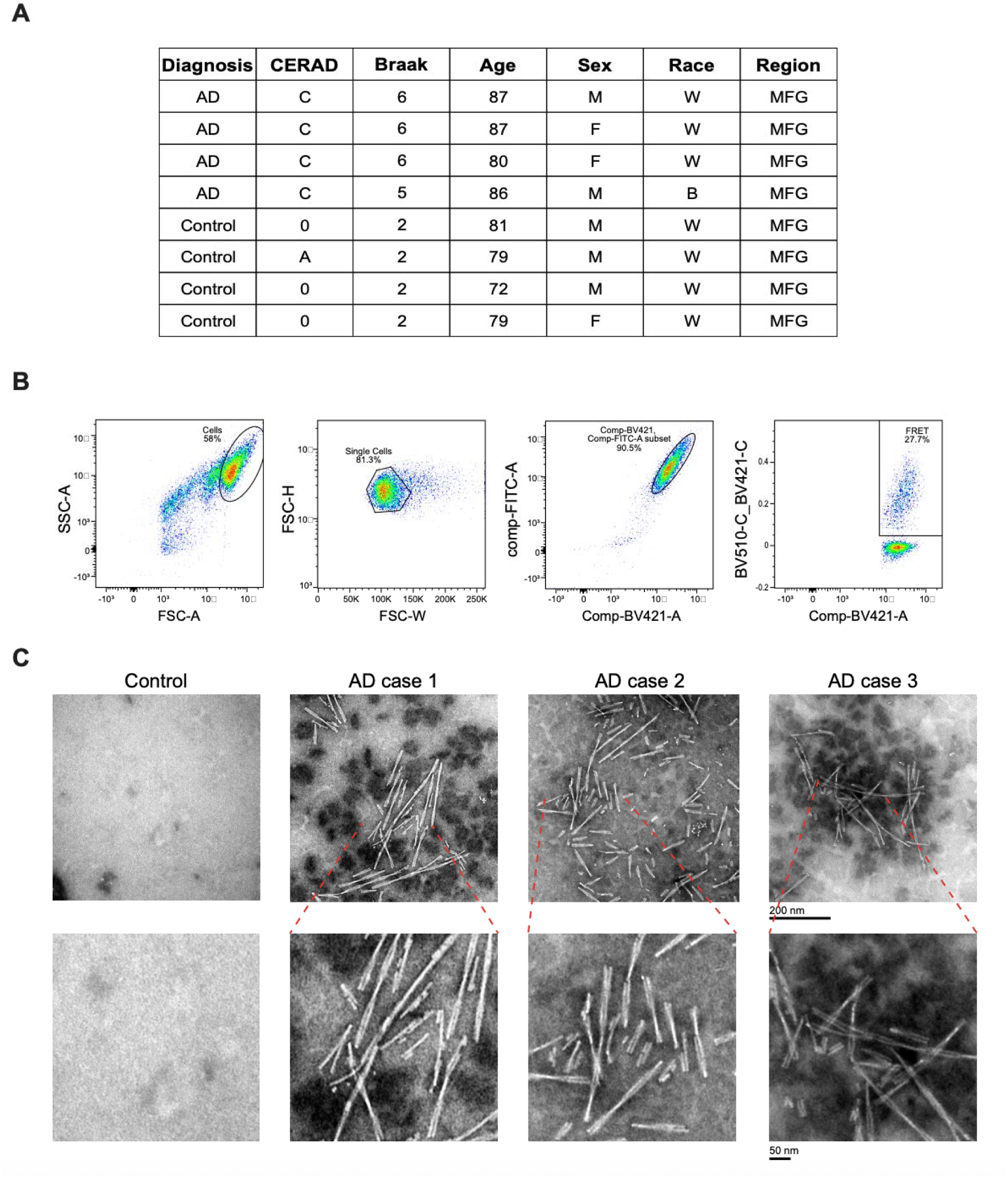
Isolation of Tau-seed from AD brain samples, related to. Figure 1 (A) Summary of human brain samples used in this study. All tissues were obtained from the middle frontal gyrus (MFG). (B) Flow cytometry gating strategy for quantifying FRET-positive cells. Cells were identified on SSC-A vs FSC-A. Single-cell populations were identified by FSC-H vs FSC-W gating. Non-fluorescent cells and cells with extreme deviations in YFP/CFP ratio were excluded on BV421-A vs FITC-A to prevent false positives. FRET-positive cells were gated on BV520-A/BV421-A ratio vs BV421-A based on elevated BV510-A fluorescence relative to BV421-A, with thresholds defined using negative controls. (C) Tau twisted filaments detected by electron microscopy (EM) in the Tau immunoprecipitation (Tau-IP) product from AD SEC fraction 9 (F9). No Tau filaments were detected in the corresponding control SEC F9. Scale bar: 200 nm; inset: 50 nm.

**Figure S2.**
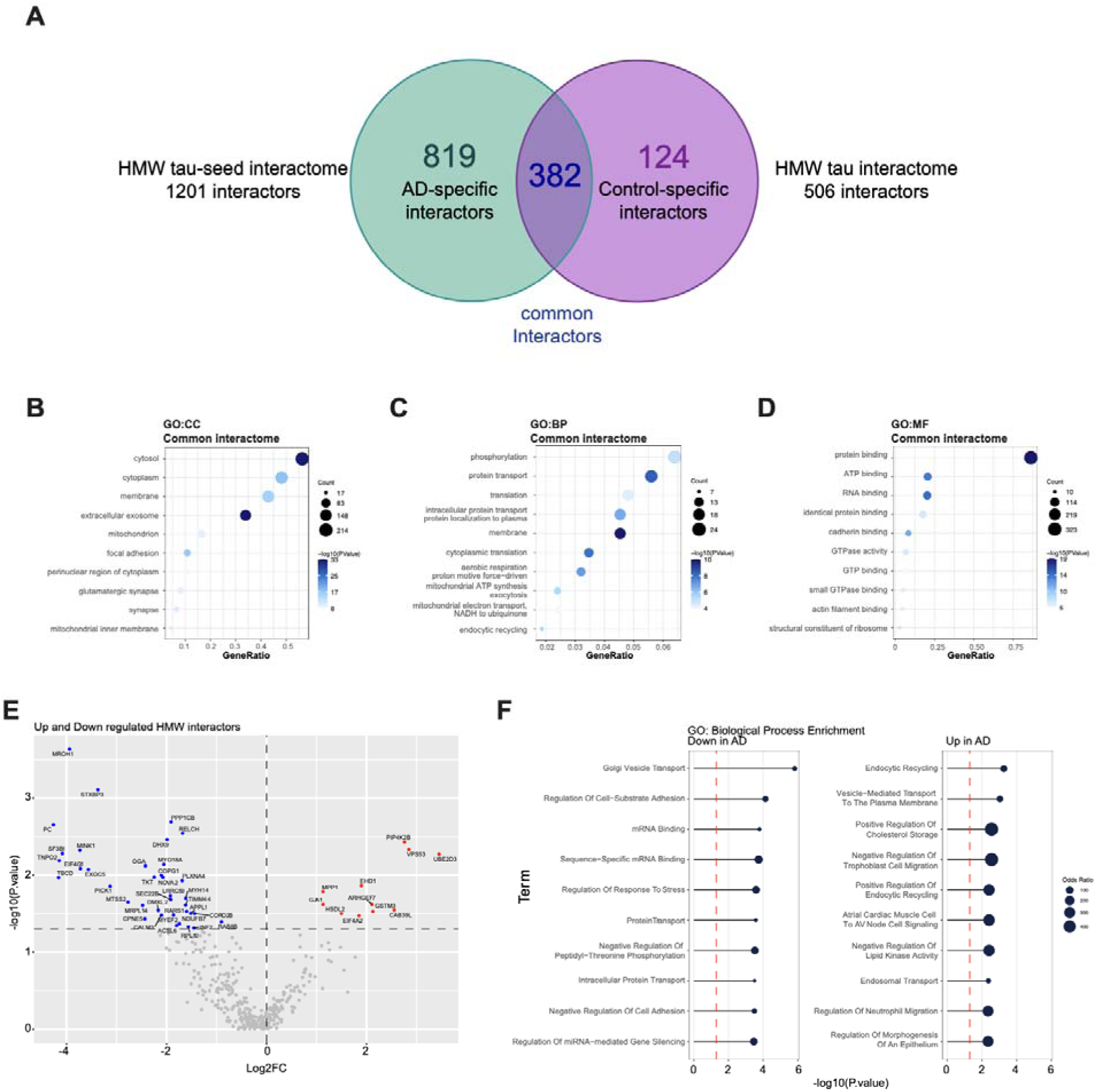
Distinct abundance profiles of shared Tau interactors in AD and control, related to Figure 2 (A) Venn analysis of interactors in AD Tau-seed and HMW-Tau datasets, highlighting 382 shared proteins. (B-D) GO enrichment of shared interactors across Cellular Component (B), Biological Process (C), and Molecular Function (D). See also Table S1. (E) Volcano plot of distinct abundances of shared Tau interactors in AD vs. control. See also Table S1. (F) GO enrichment of shared Tau interactors downregulated (F) or upregulated (G) in AD compared to control. Top 10 enriched terms are shown. See also Table S1.

**Figure S3.**
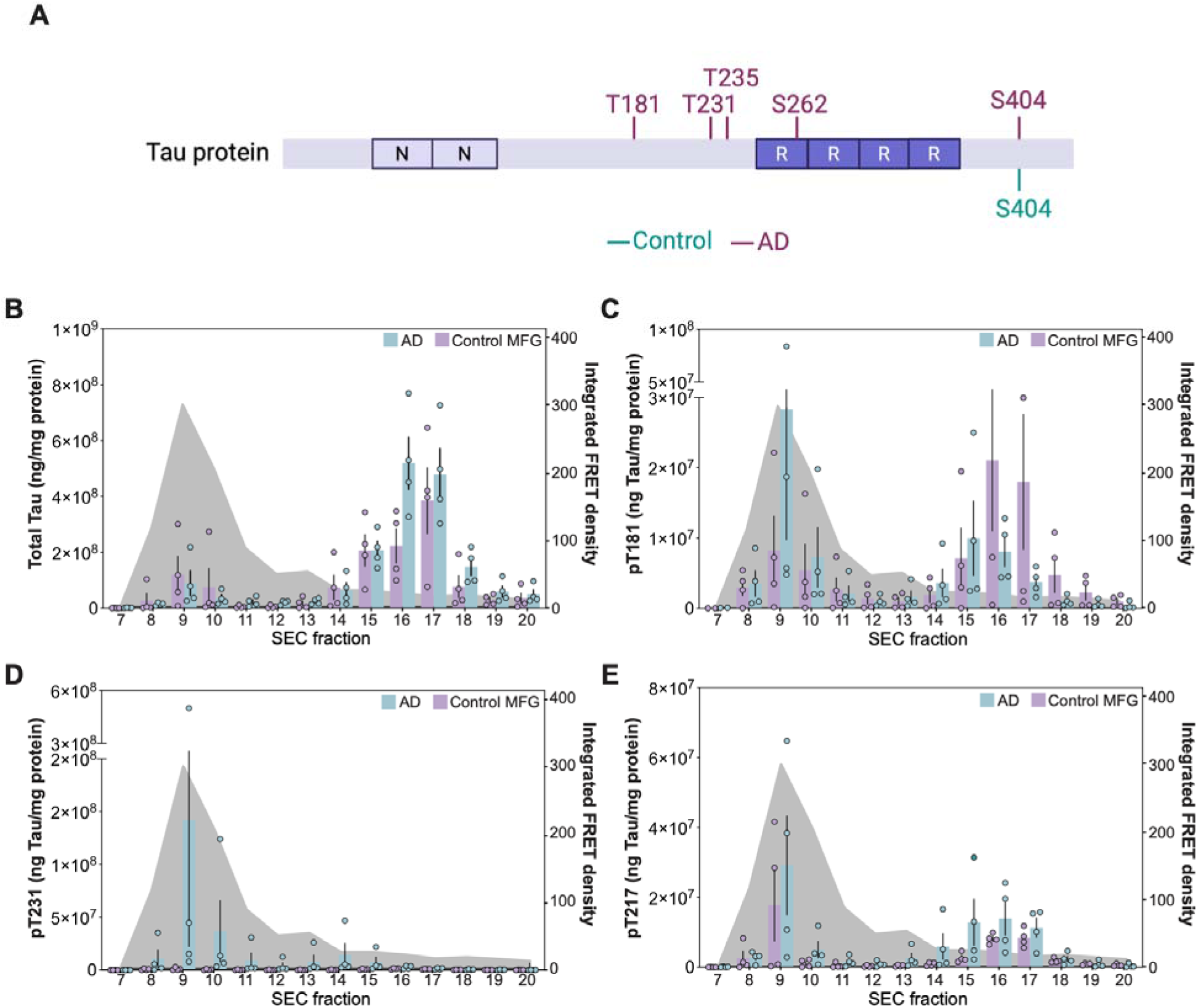
Differential phosphorylation signatures displayed by AD Tau-seed, related to Figure 2 (A) Schematic of the human Tau protein with phosphorylation sites identified by MS. Sites specific to AD Tau-seed (purple) and control HMW-Tau (green) are highlighted. (B-E) Quantification of total Tau (B), pT181 (C), pT231 (D), and pT217 (E) by MSD assay across SEC fractions from AD and control brain lysates (MFG brain region). Seeding activity of AD SEC is shown as a grey area behind each graph. Data are shown as the mean ± s.e.m. (*n*=3).

**Figure S4.**
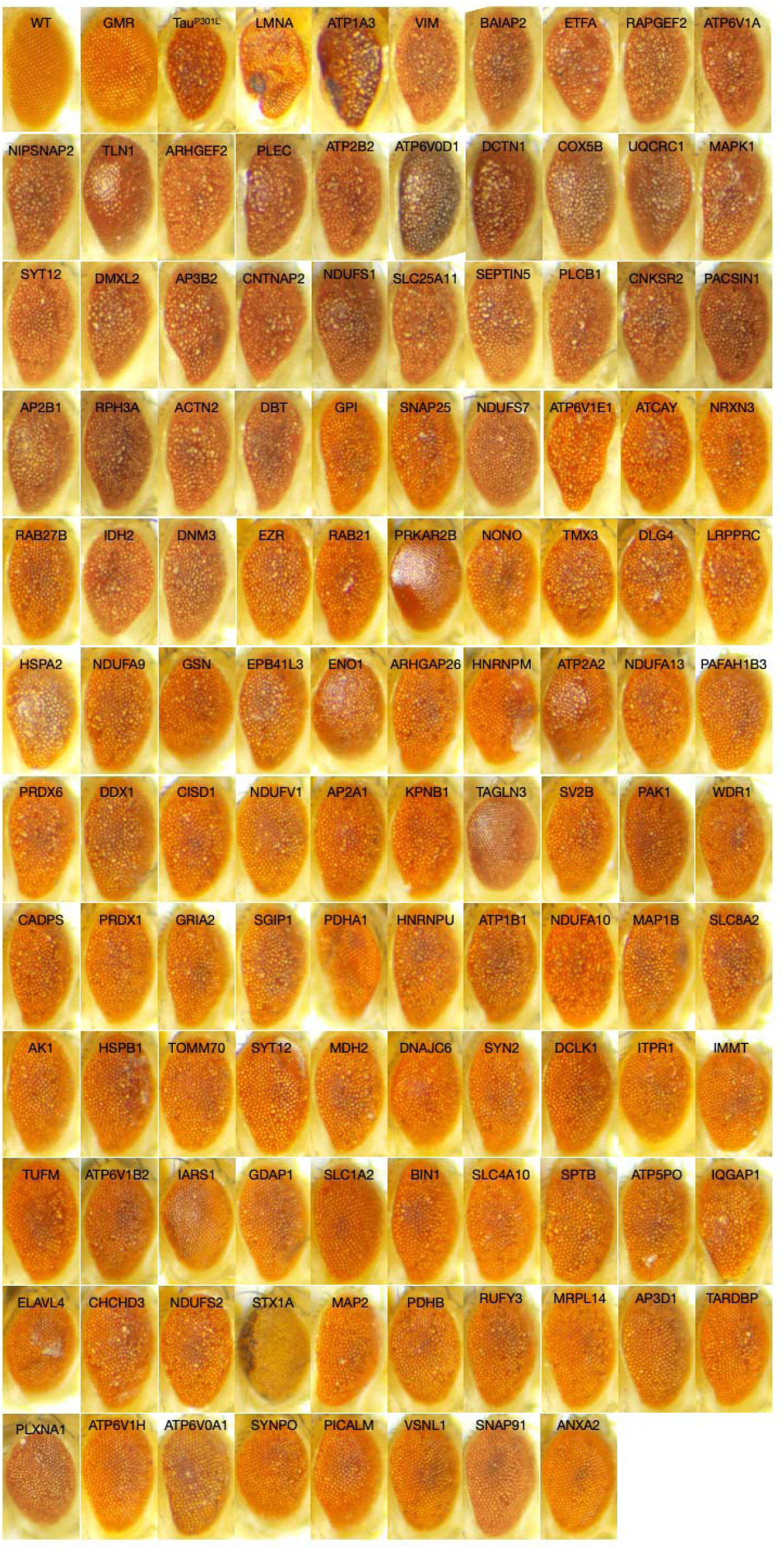
Eye phenotypes in *Drosophila* tauopathy model, related to Figure 4 (A) Representative eye images from flies expressing RNAi against 115 Tau-seed interactors.

**Figure S5.**
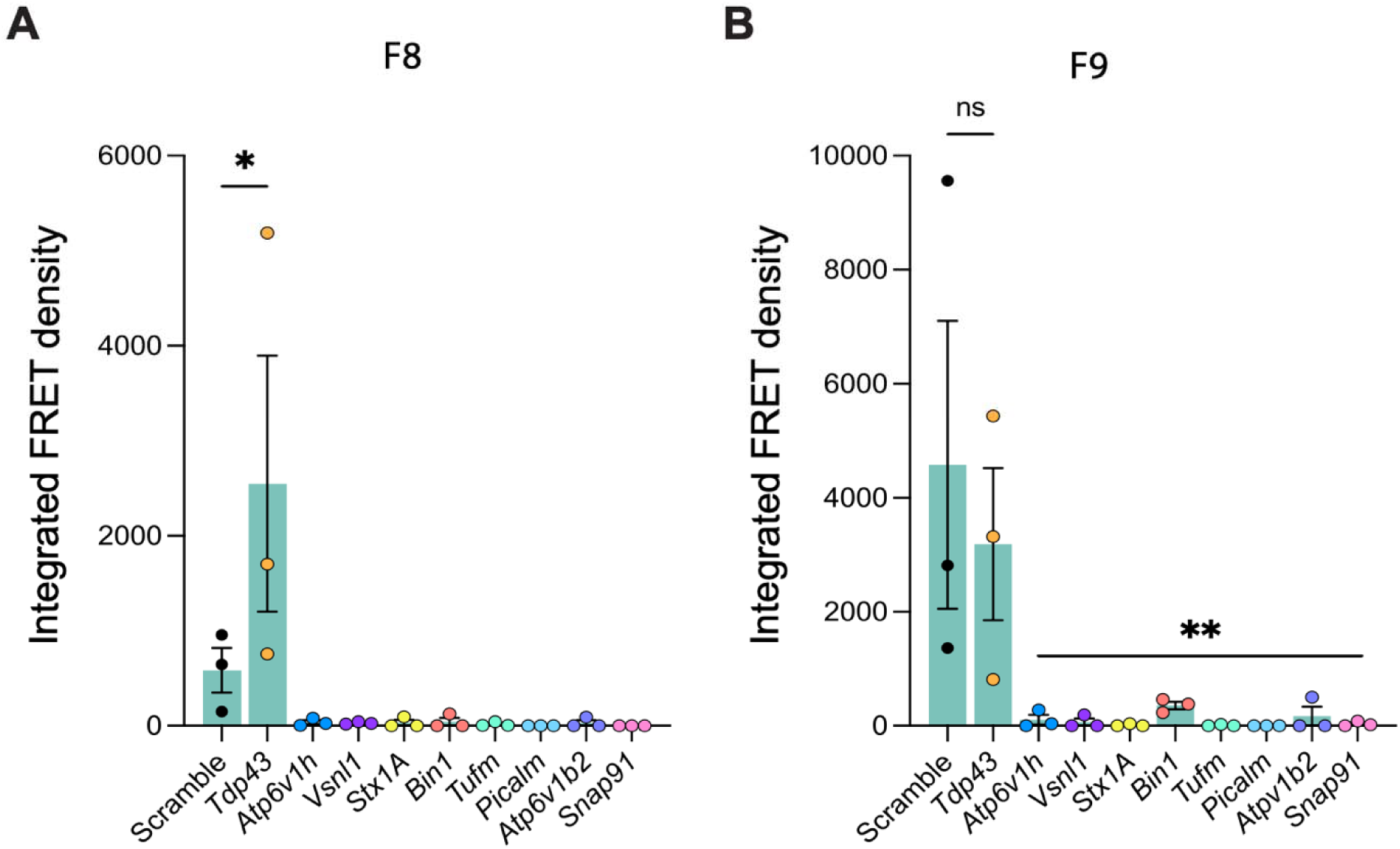
Tau seeding activity in HMW-Tau fractions following shRNA knockdown, related to Figure 6 (A and B) Tau seeding activity in SEC fractions 8 (A) and 9 (B) from PS19 brain lysates treated with shRNA targeting Tau-seed interactors. Data are shown as the mean ± s.e.m., and significance was determined by one-way ANOVA (*p<0.05, **p<0.005; *n*=3).

